## Supplementary Information for "High-parametric protein maps reveal the spatial organization in early-developing human lung"

**Supplementary Discussion Point**

A number of cell type clusters in analyses shown in ***Figure 2A-B*** and ***Figure 3A-B***, ***Figure 4***, and all related supplementary figures were not represented uniformly across all the weeks due to a number of technical reasons:

As shown by the podoplanin (PDPN) antibody staining patterns in ***Figure 1D,*** mesothelial cells were indeed present at all five developmental weeks. However, as detailed in ***Supplementary Figure 4A***, in the relatively larger tissues of the 8.5 week and all later-stage lung samples, we experienced uneven signal intensity for various markers in peripheral regions, causing issues in downstream clustering after image segmentation. Thus, we removed these peripheral regions before downstream data analysis. This impacted the representation of mesothelial cells in all weeks except the week 6 in the downstream analyses.

Chondroblasts were missing in the 12 week old sample because the section presented for the 12 week old sample originated from a relatively more peripheral region of the organ, representing more distal parts of the airway network as compared to other samples.

For the pericytes and the airway fibroblasts, we did not have single specific markers for these cell types. Instead, similar to neuronal cells, we used a combination of several markers that provided clusters with characteristic signatures aligning with the positioning and morphology of these cells in the tissue images. This approach, while demonstrating the strength of multi-parametric image analysis at the protein level, is prone to underperform if one or more of the markers do not perform equally well (*e.g.* ACTA2 in week 12 compared to weeks 11 and 13), affecting the clustering performance.

Adventitial fibroblasts are localized within the bronchovascular bundles of the lung[1]. Among the analyzed sections, only those from the week 11 sample contained such a central part of the organ. This provides an explanation about their absence in more peripheral lung sections.

**Supplementary Table 1. List of and detailed information on all evaluated antibodies.** Detailed information such as catalogue number, clone ID, target, selection reason, conjugated barcode, indirect IF dilutions and tested tissues, failed steps for all evaluated antibodies in the study. In the "Failed step" column, the step where antibodies failed is reported according to the evaluation after the indirect IF, conjugation or multiplexed imaging run.

**Supplementary Table 2. Values for background subtraction.** Values for background subtraction for all the markers and developmental weeks.

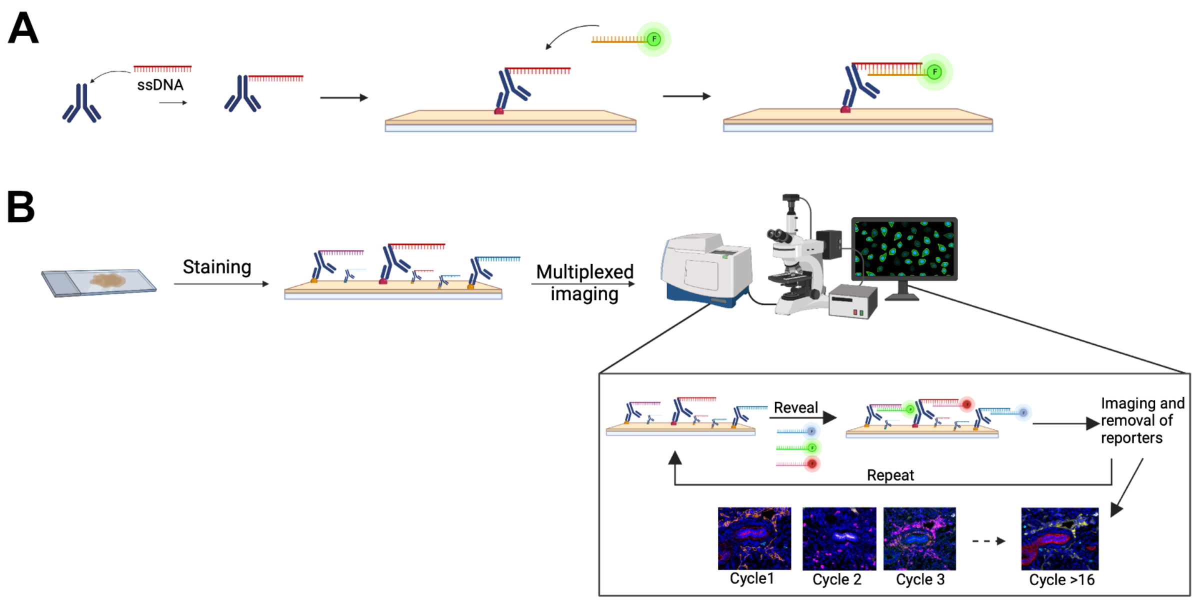


**Supplementary Figure 1. High-parametric tissue imaging workflow. A)** Steps of antibody conjugation and conjugation validation. Antibodies were conjugated with single-stranded DNA (ssDNA) and validation of the conjugated antibodies was done with reporters (complementary DNA strands conjugated to fluorophores). **B)** Staining the tissue with conjugated antibodies. During each cycle of the automated image acquisition run, up to three reporters were revealed to visualize up to three different antibodies. After each imaging cycle, the reporters were removed to reveal the next set of reporters. (Figure was created with BioRender.com.)

Week 6

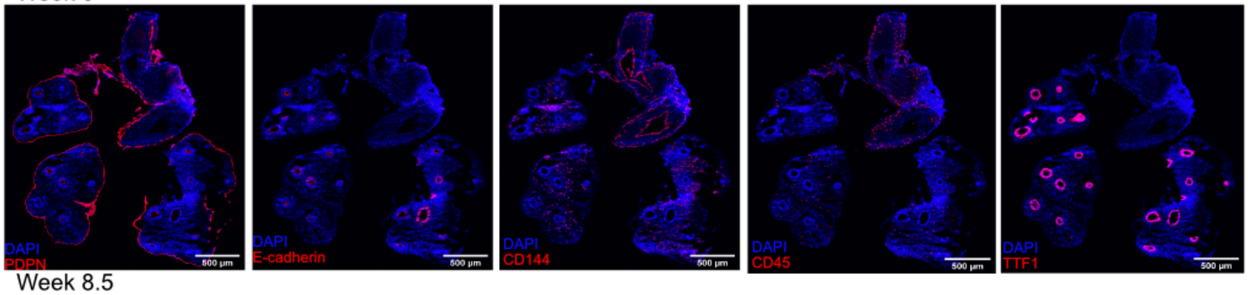


Week 8.5

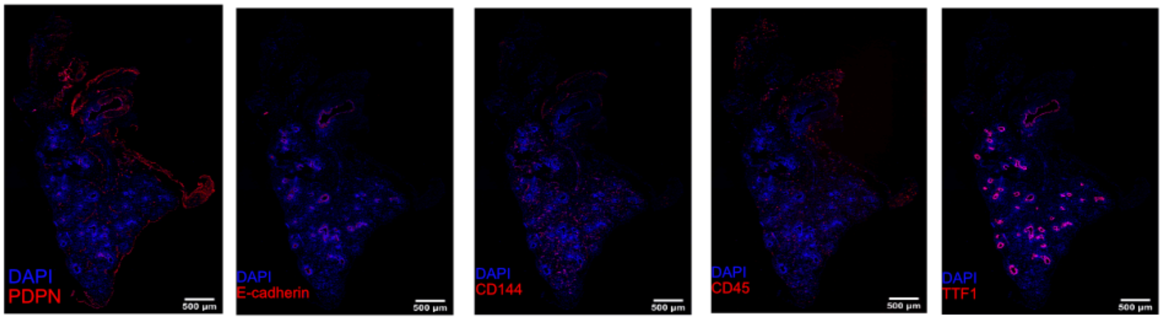


Week 11

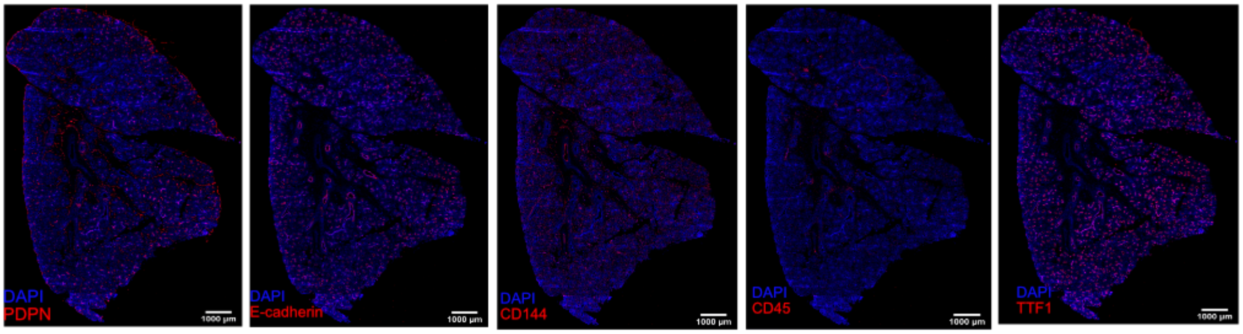


Week 12

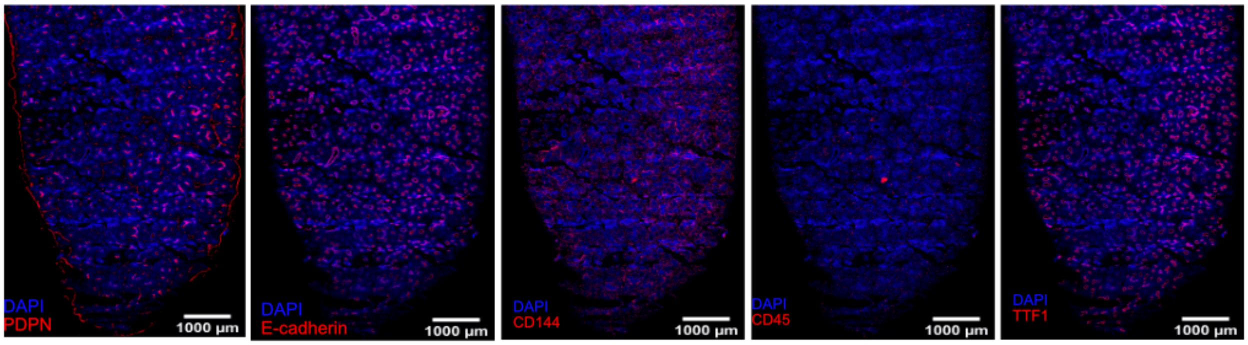


Week 13

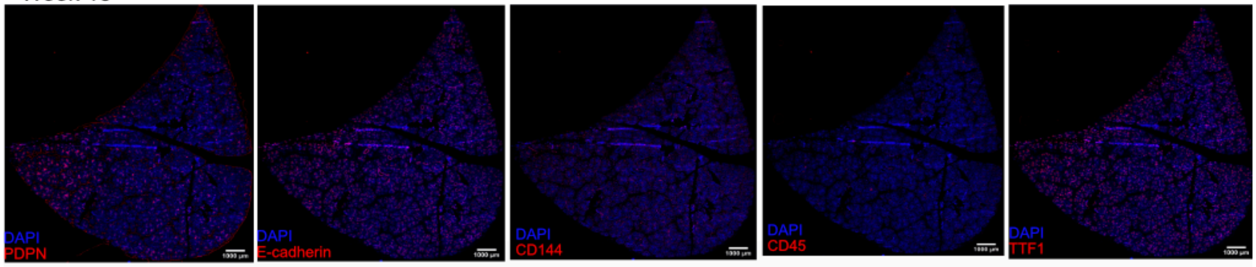

**Supplementary Figure 2. Original imaging area for the lung tissues from different developmental weeks.** Staining patterns of PDPN (lymphatic endothelial and mesothelial marker), CDH1 (E-Cadherin, epithelial marker), CD144 (endothelial marker), CD45 (general immune marker) and TTF1 (epithelial marker) are shown in red on the original imaging area. Scale bar corresponds to 500 μm in weeks 6 and 8.5, and to 1000 μm in weeks 11, 12 and 13.

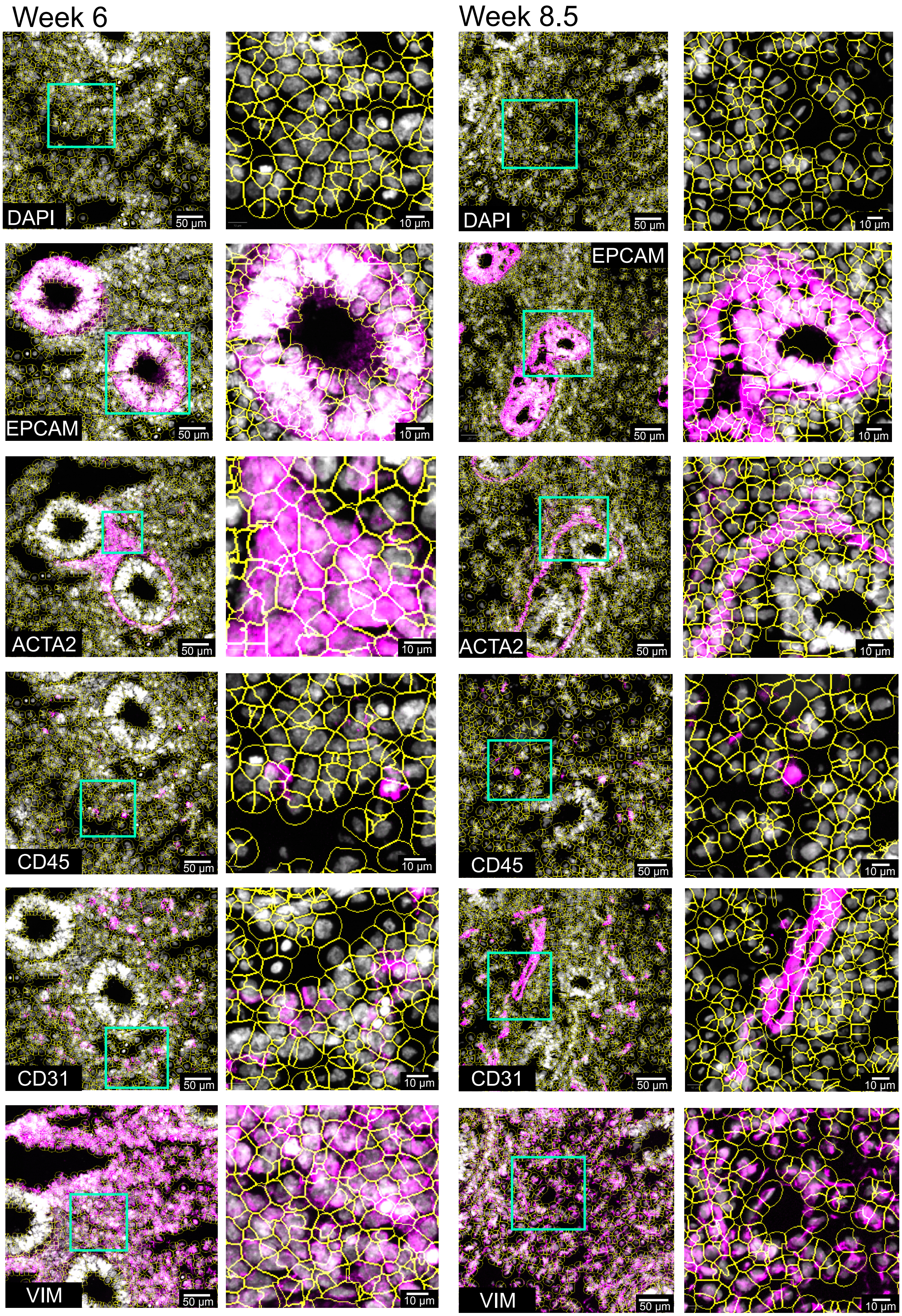
Week 6
Week 8.5
DAPI
EPCAM
ACTA2
CD45
CD31
VIM
50 µm
10 µm

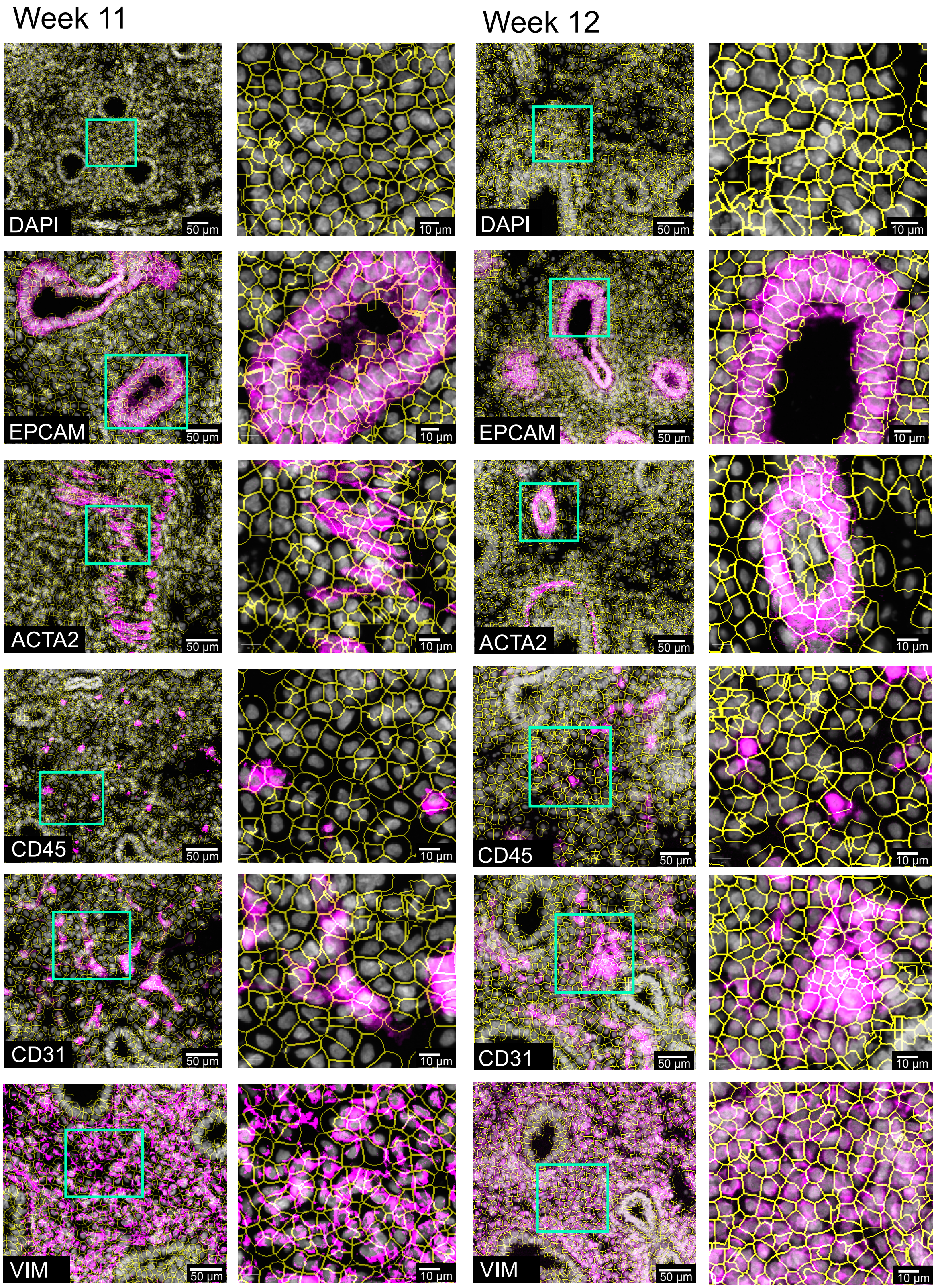

Week 11
Week 12
DAPI
EPCAM
ACTA2
CD45
CD31
VIM
50 µm
10 µm

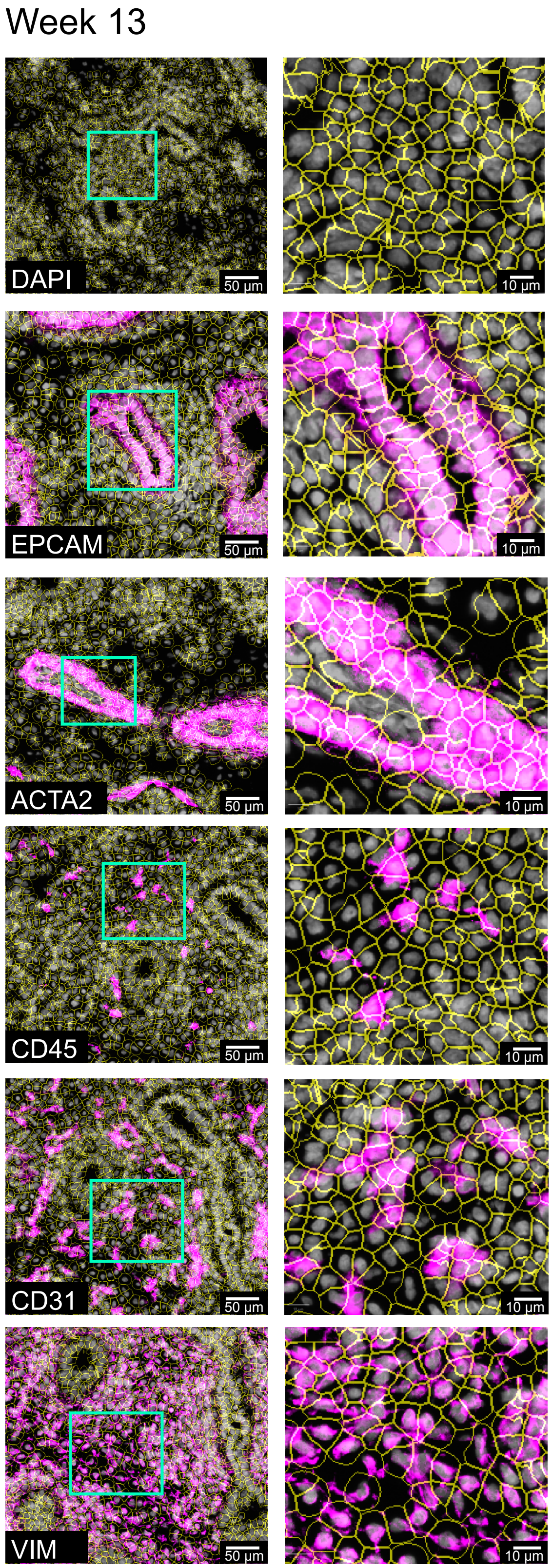


**Supplementary Figure 3. Segmentation of the developing human lung tissue imaging data.** DAPI and EPCAM images were used to create segmentation masks in PIPEX (https://github.com/CellProfiling/pipex.git) and the resulting .csv file included the intensities of each marker, cell ID and coordinates of each segmented cell. For each week, the images in the left column show selected regions to demonstrate the fit of the segmentation mask for DAPI (nuclear marker), EPCAM (epithelial cells), ACTA2 (smooth muscle cells), CD45 (immune cells), CD31 (endothelial cells) and VIM (mesenchymal cells). The images in the right column display zoomed-in views within the green inset boxes. Scale bars correspond to 50 μm in the images on the left side, and to 10 μm in the images on right side for the zoomed-in green boxes.

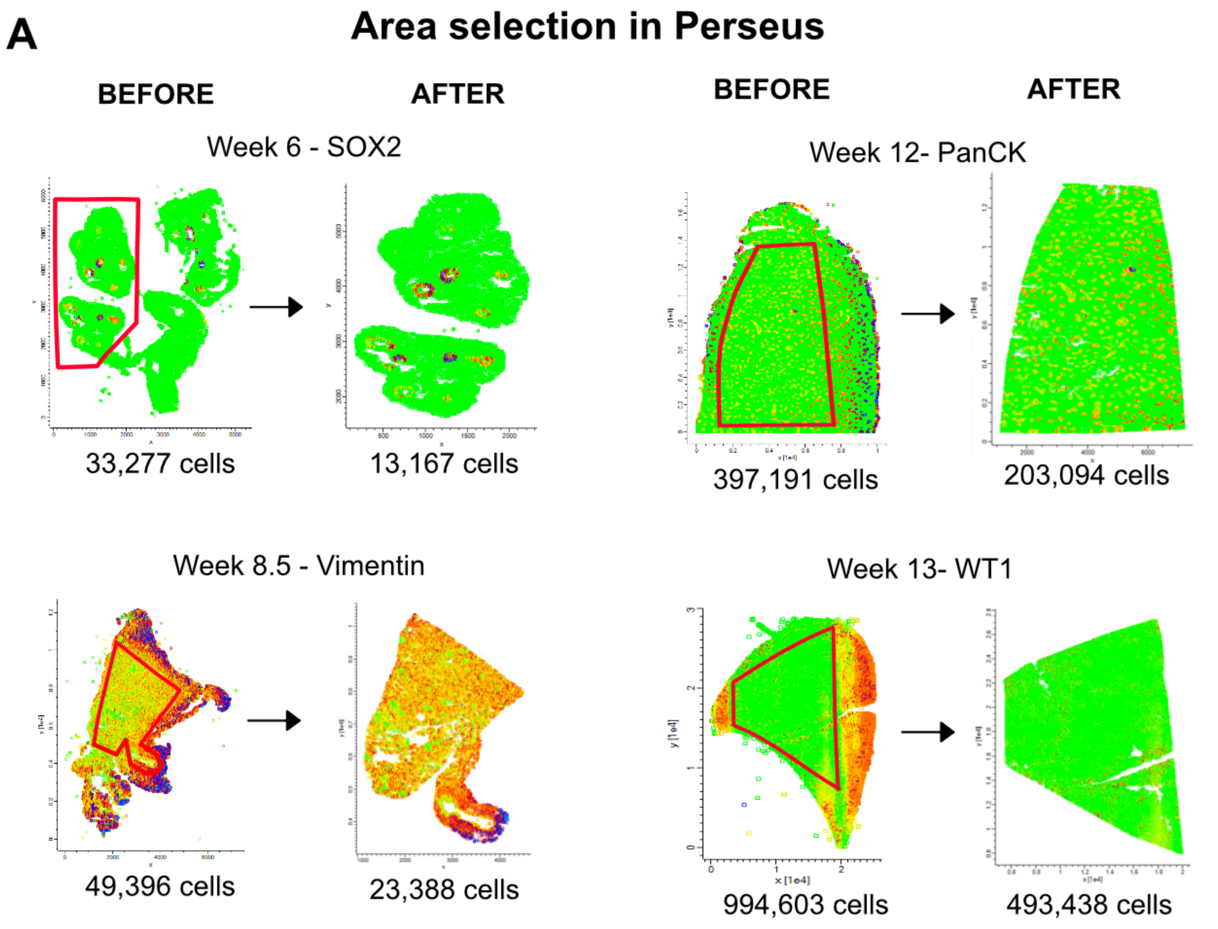


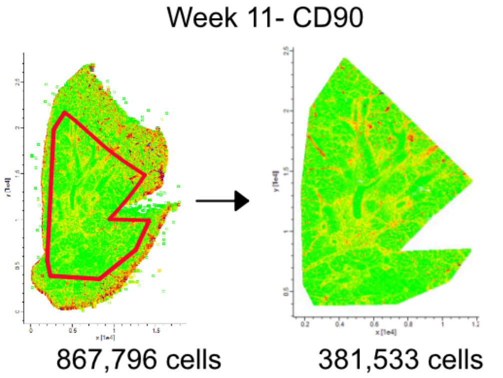


**B** Number of cells after different stages of data processing

| | 6 pcw | 8.5 pcw | 11 pcw | 12 pcw | 13 pcw |
|---|---|---|---|---|---|
| Segmentation | 34,930 | 643,84 | 978,277 | 493,588 | 1,341,495 |
| DAPI & EPCAM filtration | 33,277 | 49,396 | 867,796 | 397,191 | 994,603 |
| Area selection | 13167 | 27970 | 381,533 | 203,094 | 493,438 |
| Artifact removal | 11,713 | 24,837 | 350,208 | 179,615 | 355,198 |

**C** Number of cells per mm$^2$

| Week(pcw) | Number of cells | Area(mm$^2$) | Cells/mm$^2$ |
|---|---|---|---|
| 6 | 11,713 | 5.1 | 2,319 |
| 8.5 | 24,837 | 10.5 | 2,373 |
| 11 | 350,208 | 136.0 | 2,576 |
| 12 | 179,615 | 62.8 | 2,862 |
| 13 | 355,198 | 132.1 | 2,690 |
| | | Average | 2,564 |

**Supplementary Figure 4. Selection of developing human lung tissue regions for downstream analysis. A)** Tissues were visualized back on the tissue plane as scatterplots according to the x and y coordinates of each cell. Using Perseus[2], different markers were selected for visualization of tissue at each week to examine the areas for potential imaging artifacts. On the scatter plots, green color indicates low expression and red color indicates high expression of the corresponding markers. For week 6 and 8.5, the areas not belonging to the lung tissue were excluded. For week 11, 12 and 13, the edges of the tissue with artifacts were excluded. Red polygons indicate the regions selected in Perseus for the downstream analysis. Below each scatter plot, number of cells before and after area exclusion are indicated. **B)** Number of resulting cells after different stages of image and data processing: Segmentation, DAPI and EPCAM marker intensity filtration, area selection in Perseus and artifact removal after clustering. **C)** Calculated number of cells per unit area at different weeks, where the area corresponds to the selected polygons shown in panel **(A).**

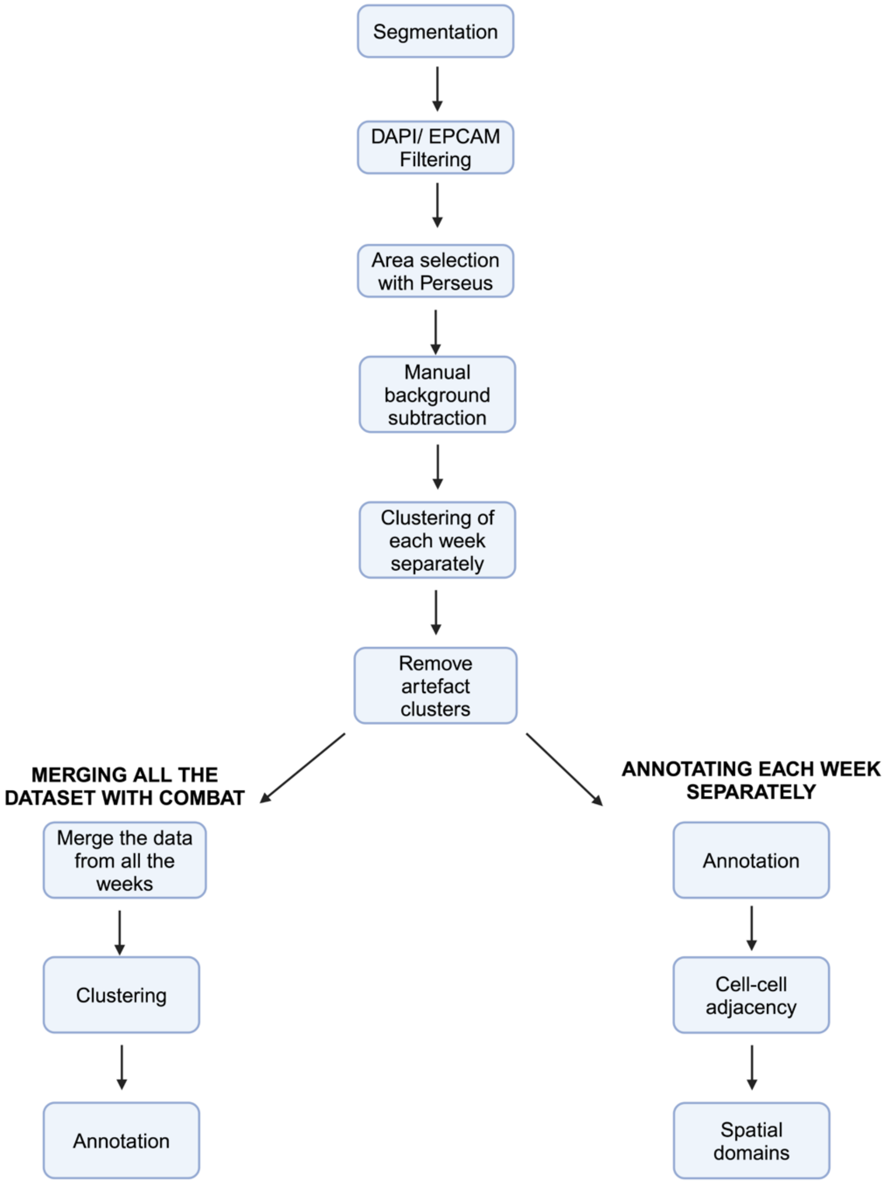


**Supplementary Figure 5. Flowchart summarizing the main steps of imaging data processing and downstream analysis.** Following image segmentation and intensity filtration for DAPI and EPCAM markers, artifact-free polygon areas were selected for each tissue using Perseus[2]. For each marker, background subtraction was performed. To identify and remove remaining artifact regions, data for each week was clustered separately and clusters annotated as artifacts were removed. Data from each week was then merged using ComBat[3] and the merged dataset was clustered and annotated to gain a global insight into the main cell types. Finally, clusters identified in each individual week were annotated for a more detailed analysis of cell types and subtypes and for a downstream analysis of cell type adjacency patterns and spatial domains in individual timepoints.

UMAP clusters before artifact removal

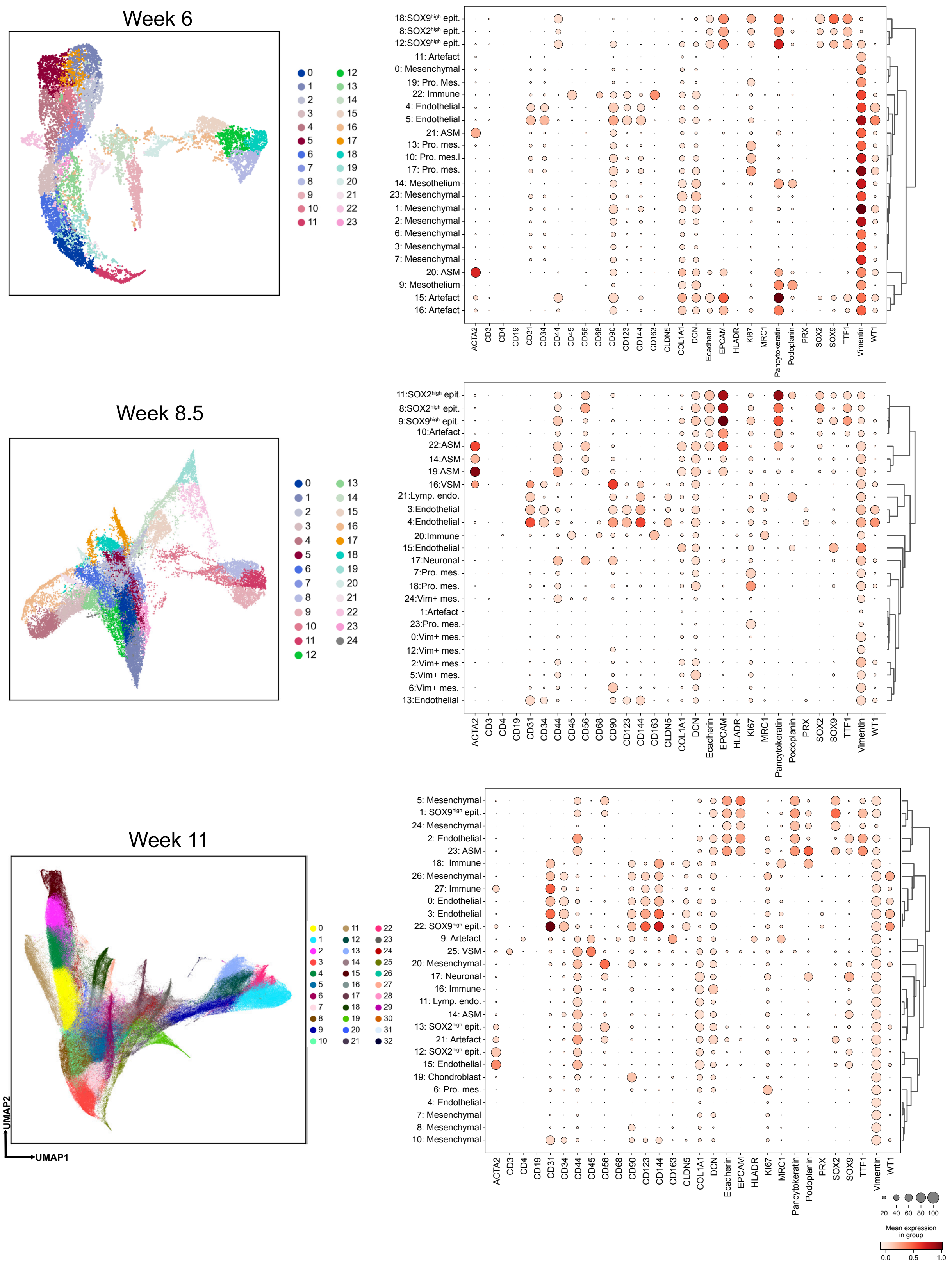

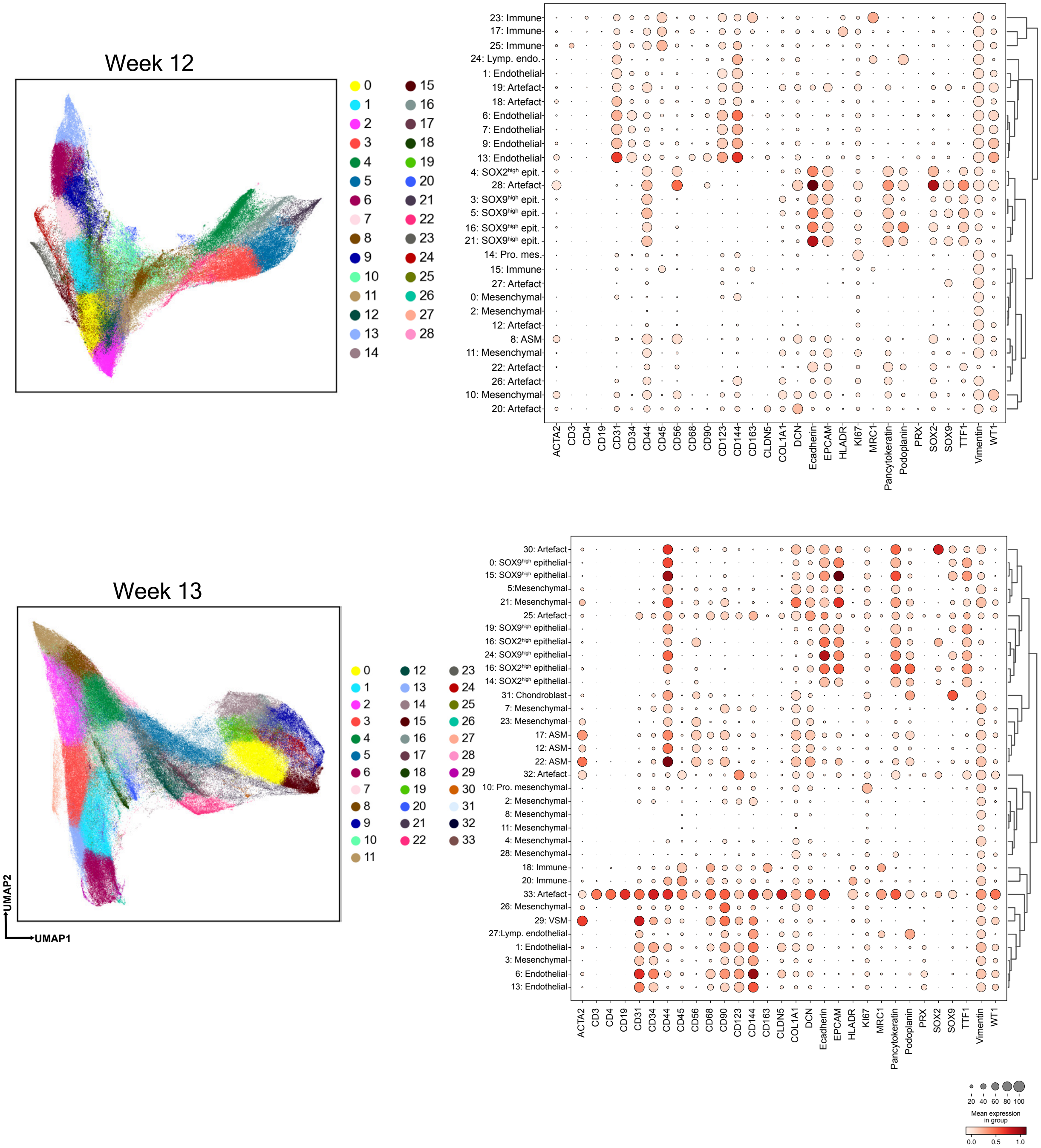


**Supplementary Figure 6. Cluster annotations for developing human lung samples at different weeks before the removal of clusters with artifacts.** UMAP plots on the left side show all clusters identified and dotplots show average marker expressions in these clusters, at each week. The size of the circles indicates the percentage of cells expressing the marker in each cluster, where dark red color indicates high and white color indicates low expression.

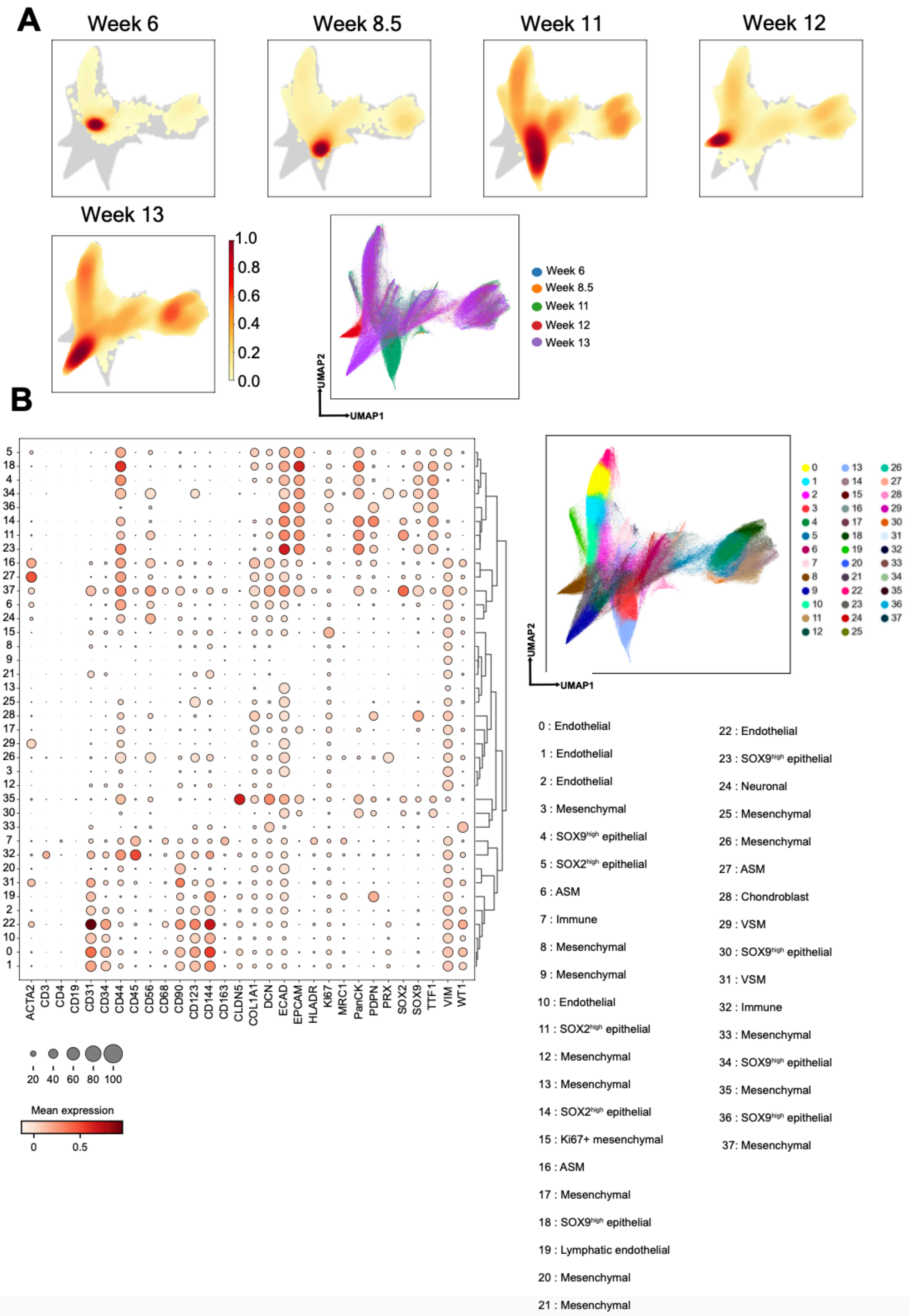
A
Week 6
Week 8.5
Week 11
Week 12
Week 13
1.0
0.8
0.6
0.4
0.2
0.0
Week 6
Week 8.5
Week 11
Week 12
Week 13
UMAP2
UMAP1
B
ACTA2 CD3 CD4 CD19 CD31 CD34 CD44 CD45 CD56 CD68 CD90 CD123 CD144 CD163 CLDN5 COL1A1 DCN ECAD EPCAM HLADR Ki67 MRC1 PanCK PDPN PRX SOX2 SOX9 TTF1 VIM WT1
20 40 60 80 100
Mean expression
0 0.5
UMAP2
UMAP1
0 : Endothelial
1 : Endothelial
2 : Endothelial
3 : Mesenchymal
4 : SOX9$^{high}$ epithelial
5 : SOX2$^{high}$ epithelial
6 : ASM
7 : Immune
8 : Mesenchymal
9 : Mesenchymal
10 : Endothelial
11 : SOX2$^{high}$ epithelial
12 : Mesenchymal
13 : Mesenchymal
14 : SOX2$^{high}$ epithelial
15 : Ki67+ mesenchymal
16 : ASM
17 : Mesenchymal
18 : SOX9$^{high}$ epithelial
19 : Lymphatic endothelial
20 : Mesenchymal
21 : Mesenchymal
22 : Endothelial
23 : SOX9$^{high}$ epithelial
24 : Neuronal
25 : Mesenchymal
26 : Mesenchymal
27 : ASM
28 : Chondroblast
29 : VSM
30 : SOX9$^{high}$ epithelial
31 : VSM
32 : Immune
33 : Mesenchymal
34 : SOX9$^{high}$ epithelial
35 : Mesenchymal
36 : SOX9$^{high}$ epithelial
37: Mesenchymal

**Supplementary Figure 7. Clustering of data merged from developing human lung samples at all analyzed weeks. A)** Density plots embedded on UMAP plots show the distribution of clusters belonging to different weeks. High density is indicated with dark red color and low intensity is indicated with light yellow color. Below, UMAP plot of the merged data is shown where blue corresponds to data points from week 6, orange to week 8.5, green to week 11, red to week 12 and purple to week 13. **B)** Dotplot in the left panel shows the average marker expression in merged data. The size of the circles indicates the percentage of cells expressing the marker in each cluster, where dark red color indicates high and white color indicates low expression. In the lower panel, annotations of each cluster of the merged data are provided.

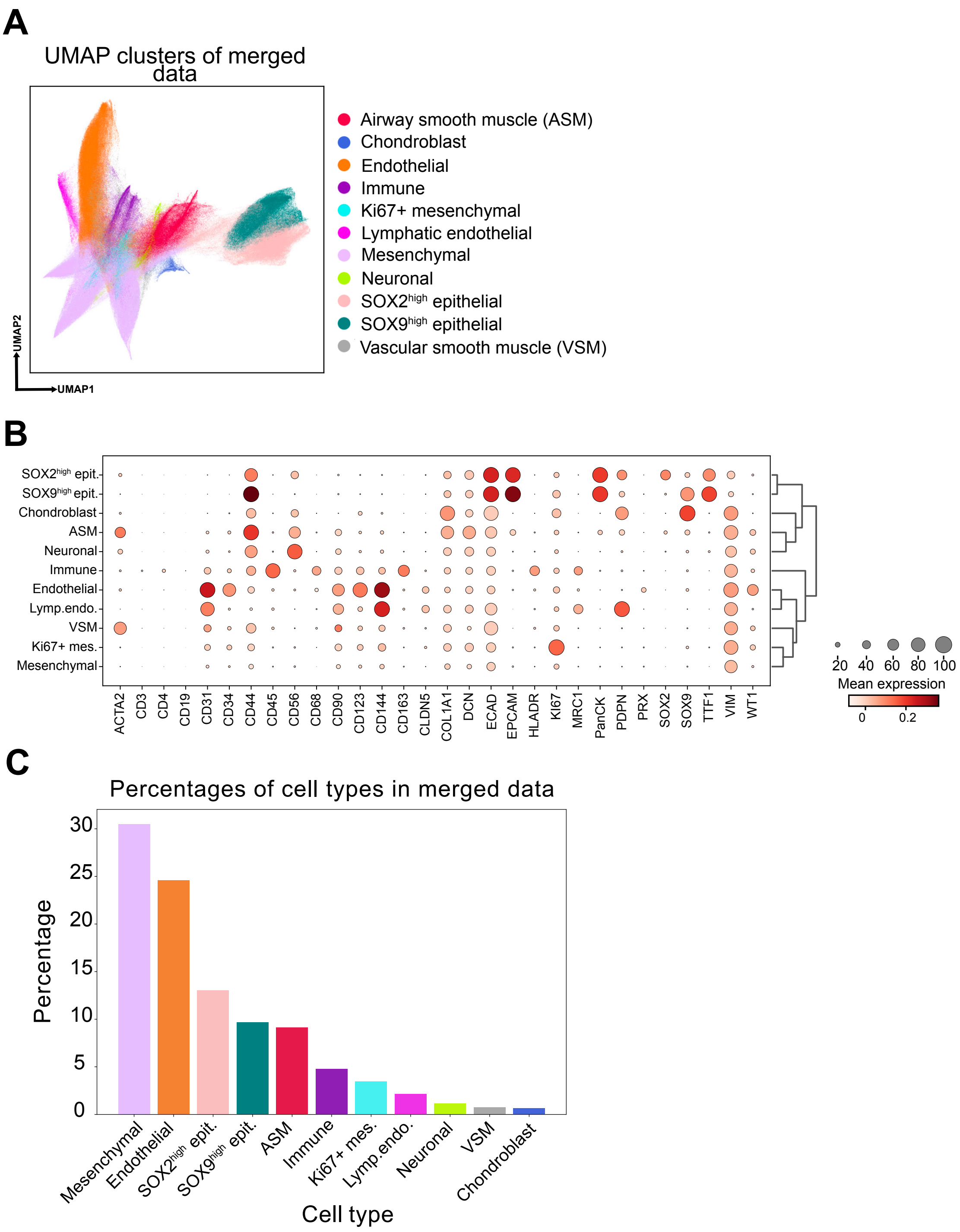


**Supplementary Figure 8. Clustering of merged human lung data across all developmental weeks. A)** Annotated cell type clusters visualized on the UMAP plot. The x-axis shows UMAP1 and the y-axis shows UMAP2. **B)** Dot plot shows the average marker expression across annotated cell type clusters. The size of the circles indicates the fraction of cells expressing the marker in each cluster. Dark red color indicates high and white color indicates low expression. **C)** Percentage of different cell types annotated in the merged data. Source data for this figure is provided within the Source Data file.

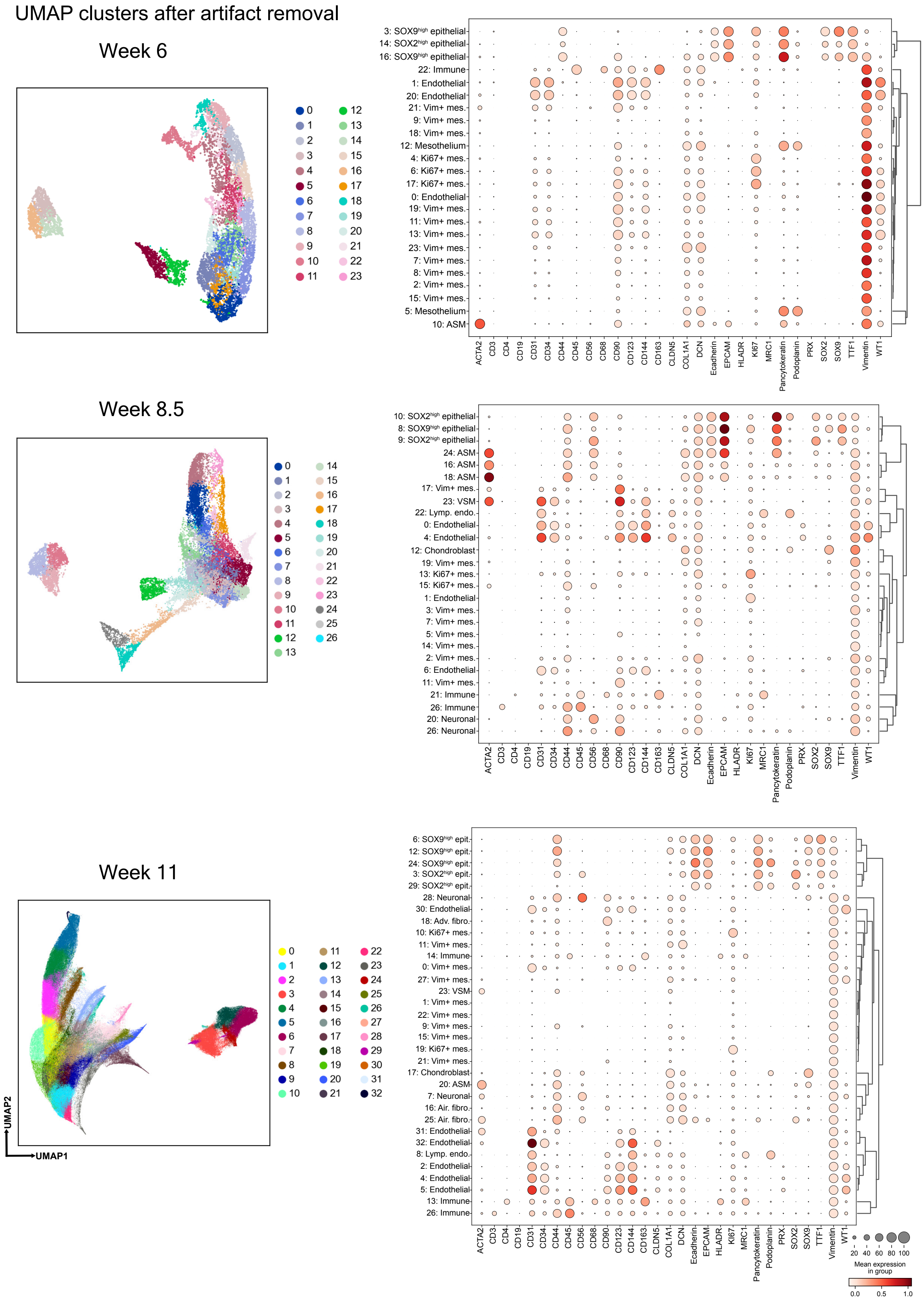

UMAP clusters after artifact removal
Week 6
Week 8.5
Week 11
UMAP2
UMAP1
3: SOX9high epithelial
14: SOX2high epithelial
16: SOX9high epithelial
22: Immune
1: Endothelial
20: Endothelial
12: Mesothelium
5: Mesothelium
10: ASM
ACTA2 CD3 CD4 CD19 CD31 CD34 CD44 CD45 CD56 CD68 CD90 CD123 CD144 CD163 CLDN5 COL1A1 DCN Ecadherin EPCAM HLADR Ki67 MRC1 Pancytokeratin Podoplanin PRX SOX2 SOX9 TTF1 Vimentin WT1
10: SOX2high epithelial
8: SOX9high epithelial
9: SOX2high epithelial
24: ASM
16: ASM
18: ASM
23: VSM
22: Lymp. endo.
12: Chondroblast
21: Immune
26: Immune
20: Neuronal
26: Neuronal
6: SOX9high epit.
12: SOX9high epit.
24: SOX9high epit.
3: SOX2high epit.
29: SOX2high epit.
28: Neuronal
18: Adv. fibro.
17: Chondroblast
20: ASM
7: Neuronal
16: Air. fibro.
25: Air. fibro.
8: Lymp. endo.
Mean expression in group

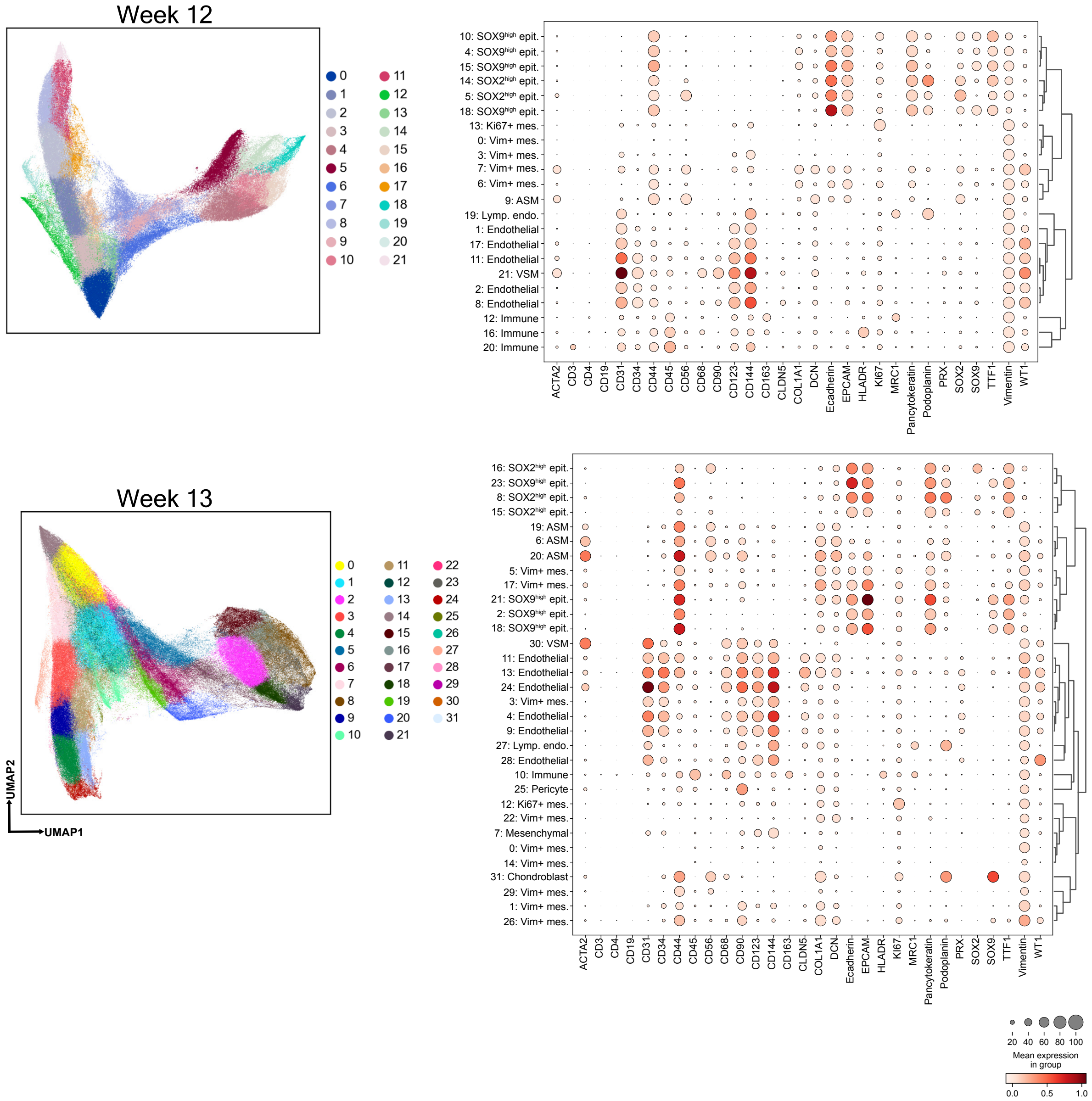


**Supplementary Figure 9. Cluster annotations for developing human lung samples at different weeks after removal of clusters with artifacts.** On the left panel, all the clusters after artifact removal are shown on the UMAP plots at each week. Dotplots to the right show average marker expression in each cluster. The size of the circles indicates the fraction of cells expressing the marker in each cluster, where dark red color indicates high and white color indicates low expression.

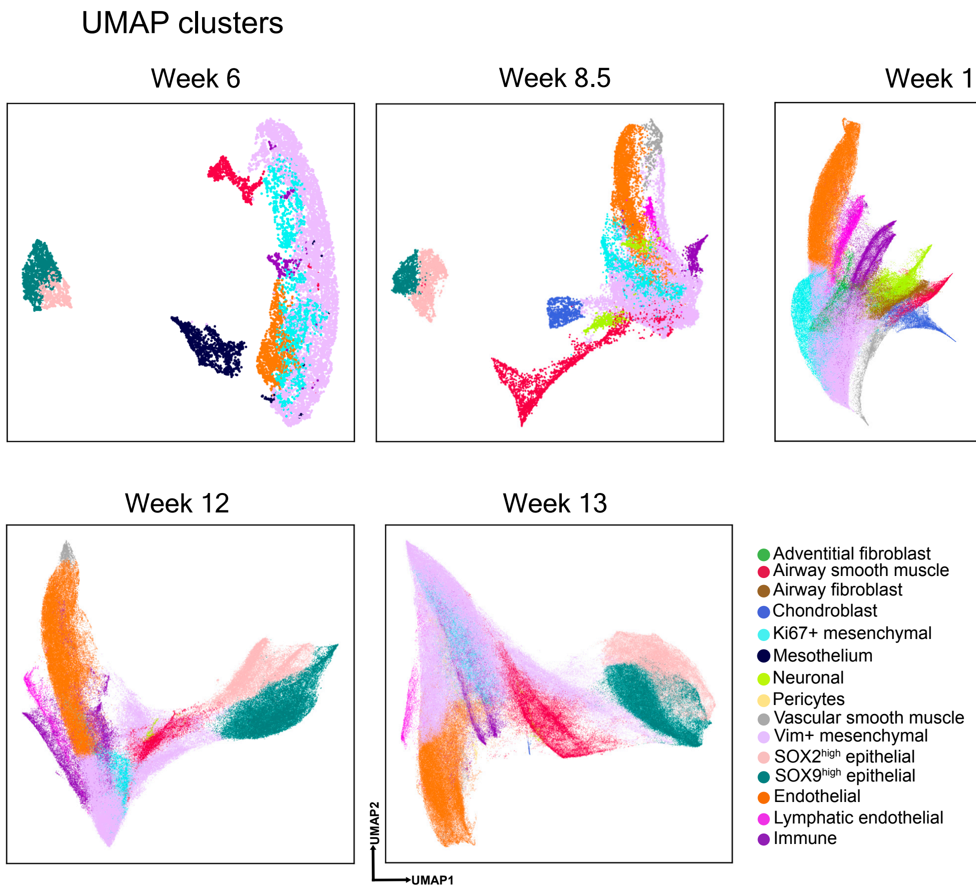


**Supplementary Figure 10. Cluster annotations for developing human lung samples at different weeks.** Clusters belonging to the same cell type were merged and plotted on UMAP embedding. Annotations of all identified clusters, represented with different colors, are provided.

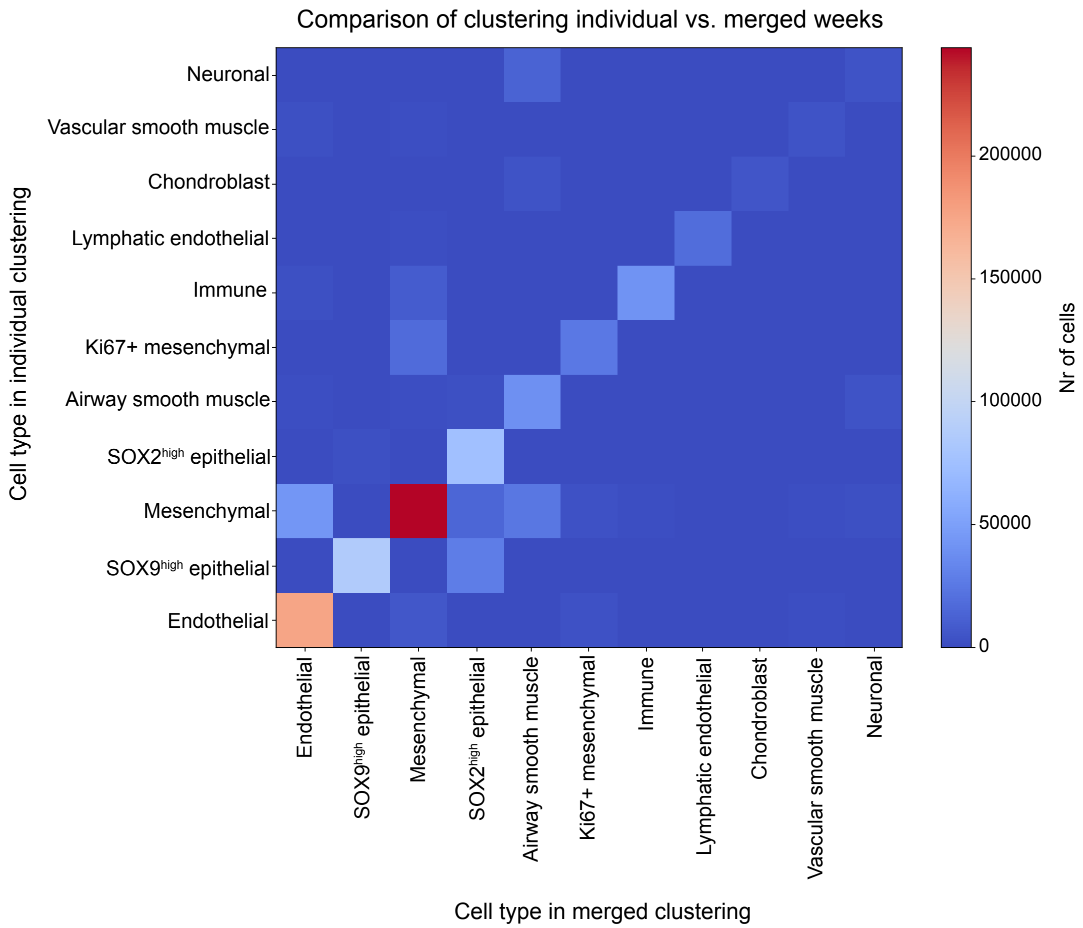


**Supplementary Figure 11. Comparison of the assigned cell type annotations in developing human lung tissue as the outcome of clustering individual versus merged timepoints.** The heatmap summarizes the count of cells assigned to one of the eleven cell types for individual and merged clustering of timepoints. While the majority (78%) of the cell type assignments were consistent, discrepancies existed in the assignment of certain cell types such as SOX2$^{high}$ epithelial cells, immune cells and airway smooth muscle cells.

**Endothelial and lymphatic endothelial structures**

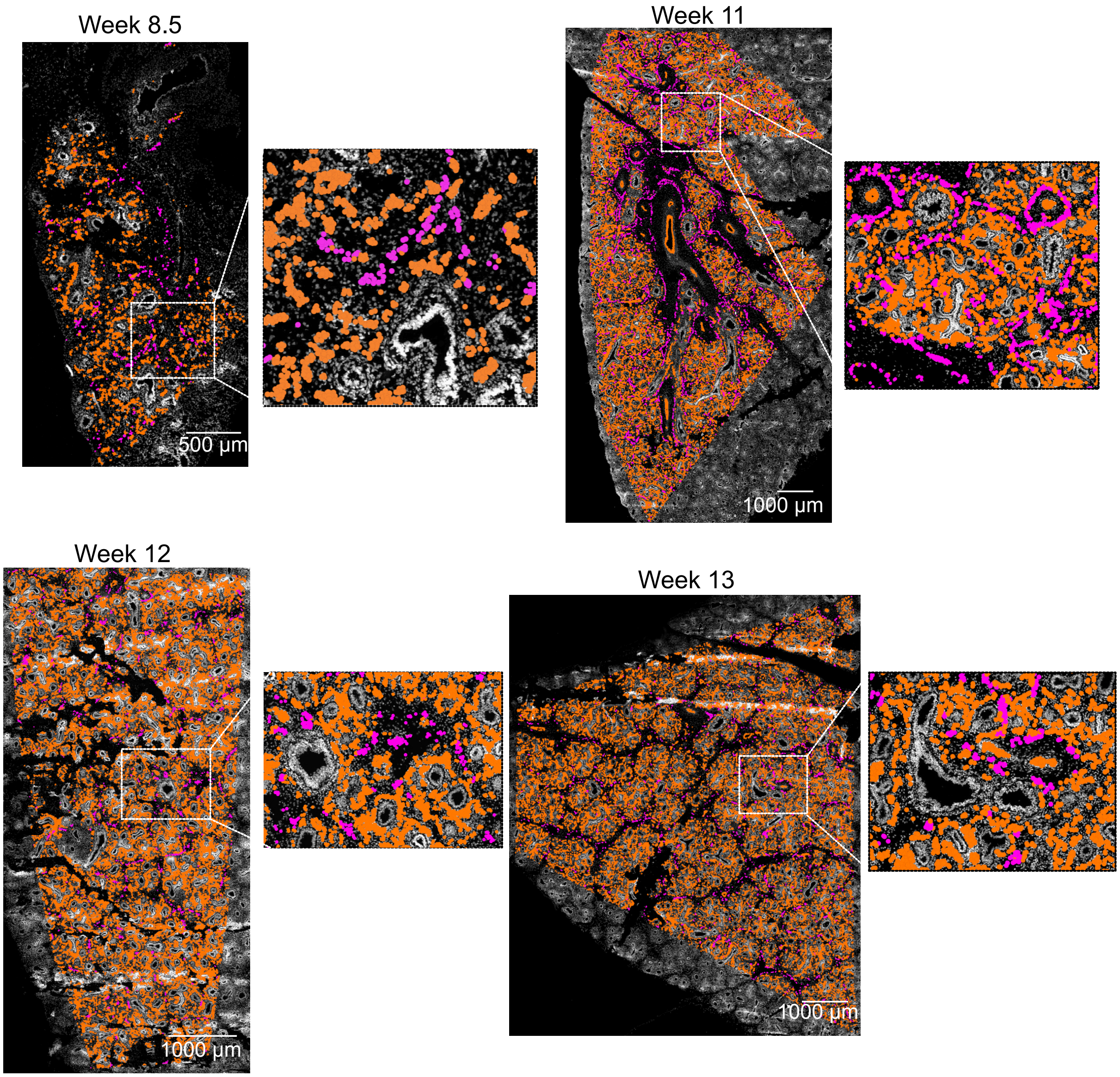


**Supplementary Figure 12. Endothelial and lymphatic endothelial structures in developing human lung.** Clusters annotated as "endothelial" (dark orange) and "lymphatic endothelial "(magenta) overlaid on DAPI channel images for each week. Scale bars correspond to 500 µm for week 8.5, and 1,000 µm for weeks 11,12 and 13.

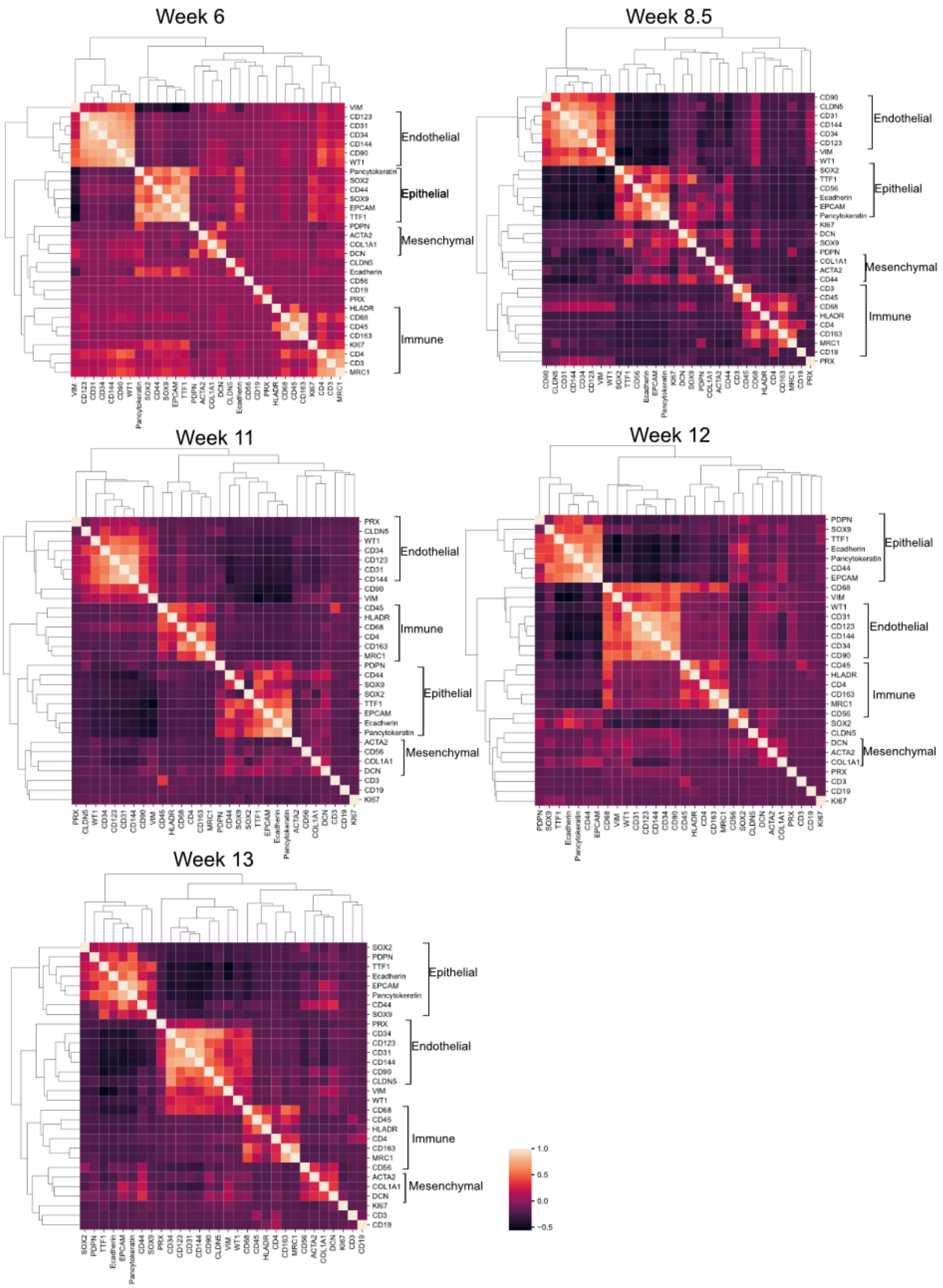
Week 6
VIM CD123 CD31 CD34 CD144 CD90 WT1 Pancytokeratin SOX2 CD44 SOX9 EPCAM TTF1 PDPN ACTA2 COL1A1 DCN CLDN5 Ecadherin CD56 CD19 PRX HLADR CD68 CD45 CD163 KI67 CD4 CD3 MRC1
Endothelial
Epithelial
Mesenchymal
Immune
Week 8.5
CD90 CLDN5 CD31 CD144 CD34 CD123 VIM WT1 SOX2 TTF1 CD56 Ecadherin EPCAM Pancytokeratin KI67 DCN SOX9 PDPN COL1A1 ACTA2 CD44 CD3 CD45 CD68 HLADR CD4 CD163 MRC1 CD19 PRX
Endothelial
Epithelial
Mesenchymal
Immune
Week 11
PRX CLDN5 WT1 CD34 CD123 CD31 CD144 CD90 VIM CD45 HLADR CD68 CD4 CD163 MRC1 PDPN CD44 SOX9 SOX2 TTF1 EPCAM Ecadherin Pancytokeratin ACTA2 CD56 COL1A1 DCN CD3 CD19 KI67
Endothelial
Immune
Epithelial
Mesenchymal
Week 12
PDPN SOX9 TTF1 Ecadherin Pancytokeratin CD44 EPCAM CD68 VIM WT1 CD31 CD123 CD144 CD34 CD90 CD45 HLADR CD4 CD163 MRC1 CD56 SOX2 CLDN5 DCN ACTA2 COL1A1 PRX CD3 CD19 KI67
Epithelial
Endothelial
Immune
Mesenchymal
Week 13
SOX2 PDPN TTF1 Ecadherin EPCAM Pancytokeratin CD44 SOX9 PRX CD34 CD123 CD31 CD144 CD90 CLDN5 VIM WT1 CD68 CD45 HLADR CD4 CD163 MRC1 CD56 ACTA2 COL1A1 DCN KI67 CD3 CD19
Epithelial
Endothelial
Immune
Mesenchymal
1.0
0.5
0.0
-0.5

**Supplementary Figure 13. Marker correlations in developing human lung samples at different weeks.** Pearson correlation coefficients are plotted in heatmaps, where Pearson's r=1 corresponds to white and r=-0.5 corresponds to black.

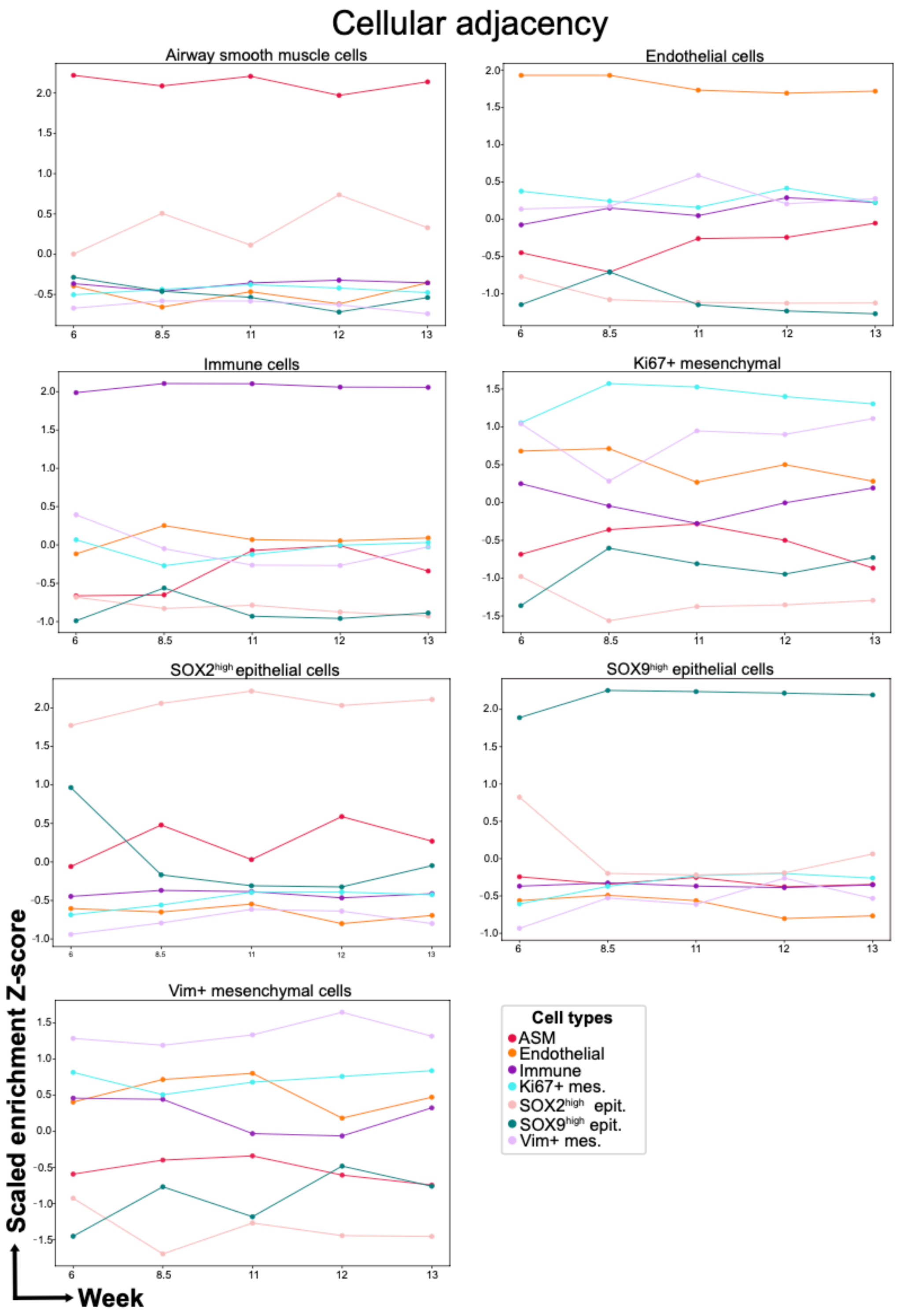


**Supplementary Figure 14. Temporal changes of scaled neighborhood enrichment Z-scores for the main cell types of developing human lung.** Each lineplot summarizes the temporal change of scaled enrichment Z-score for each main cell type for neighboring their own, or the rest of the other cell types, where each line represents one of the seven main cell types in developing human lung. Source data for this figure is provided within the Source Data file.

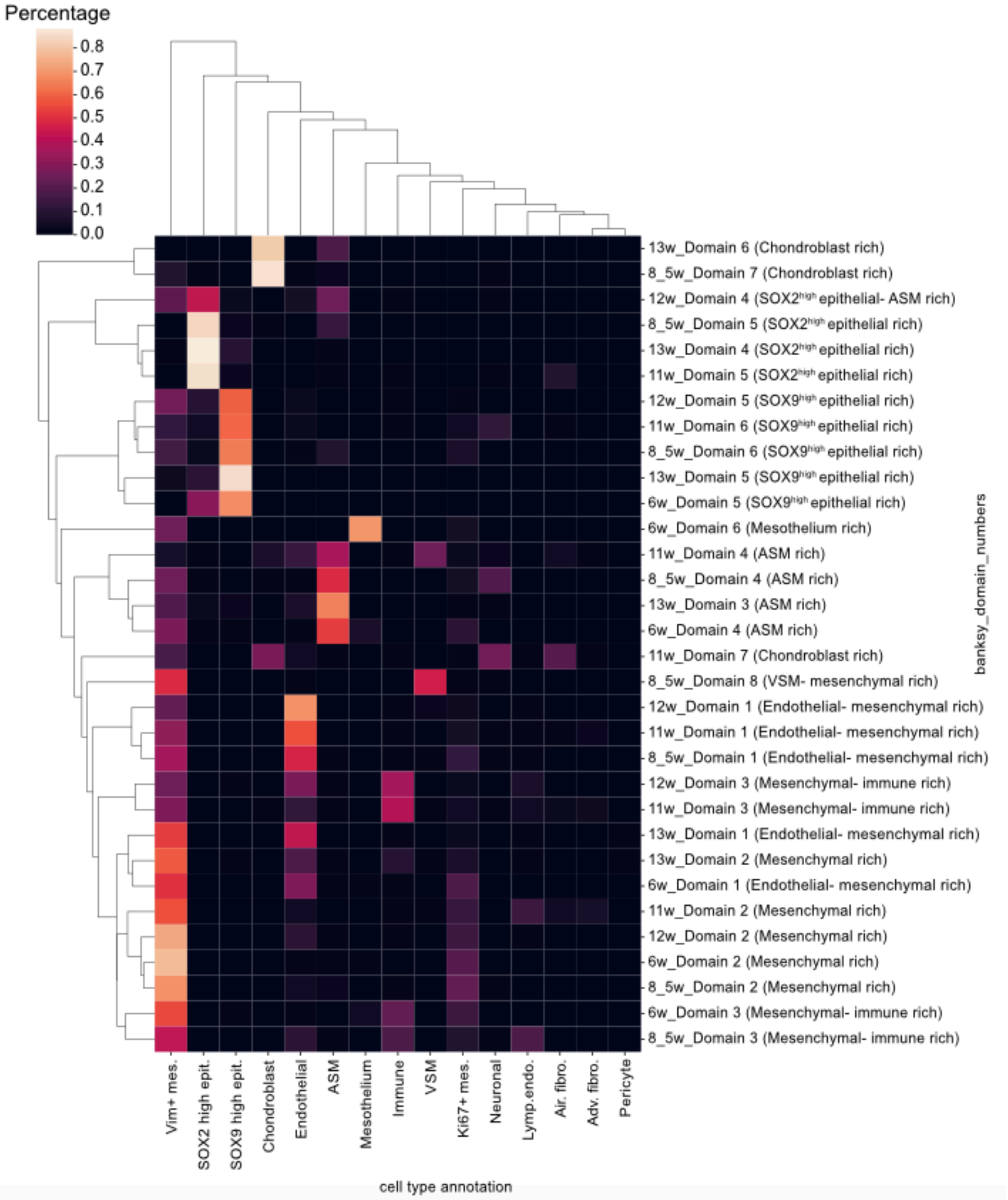


**Supplementary Figure 15. Hierarchical clustering of the cell type composition across all individual spatial domains identified in the developing human lung at different weeks.** The color scale indicates the percentage of each cell type within a domain.

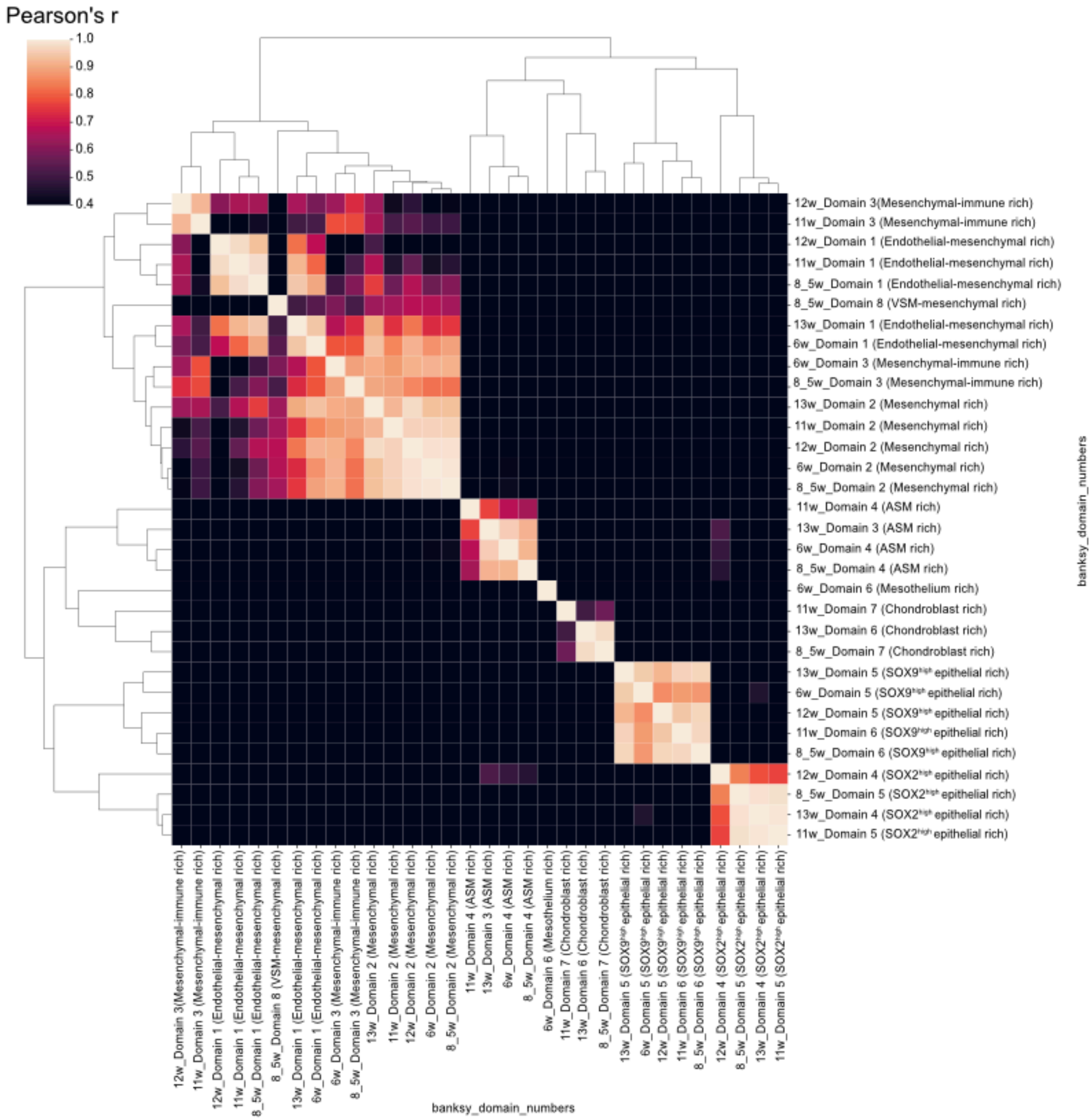


**Supplementary Figure 16. Hierarchical clustering of the correlation among all the individual spatial domains identified in the developing human lung at different weeks.** The color scale indicates Pearson correlation coefficient r.

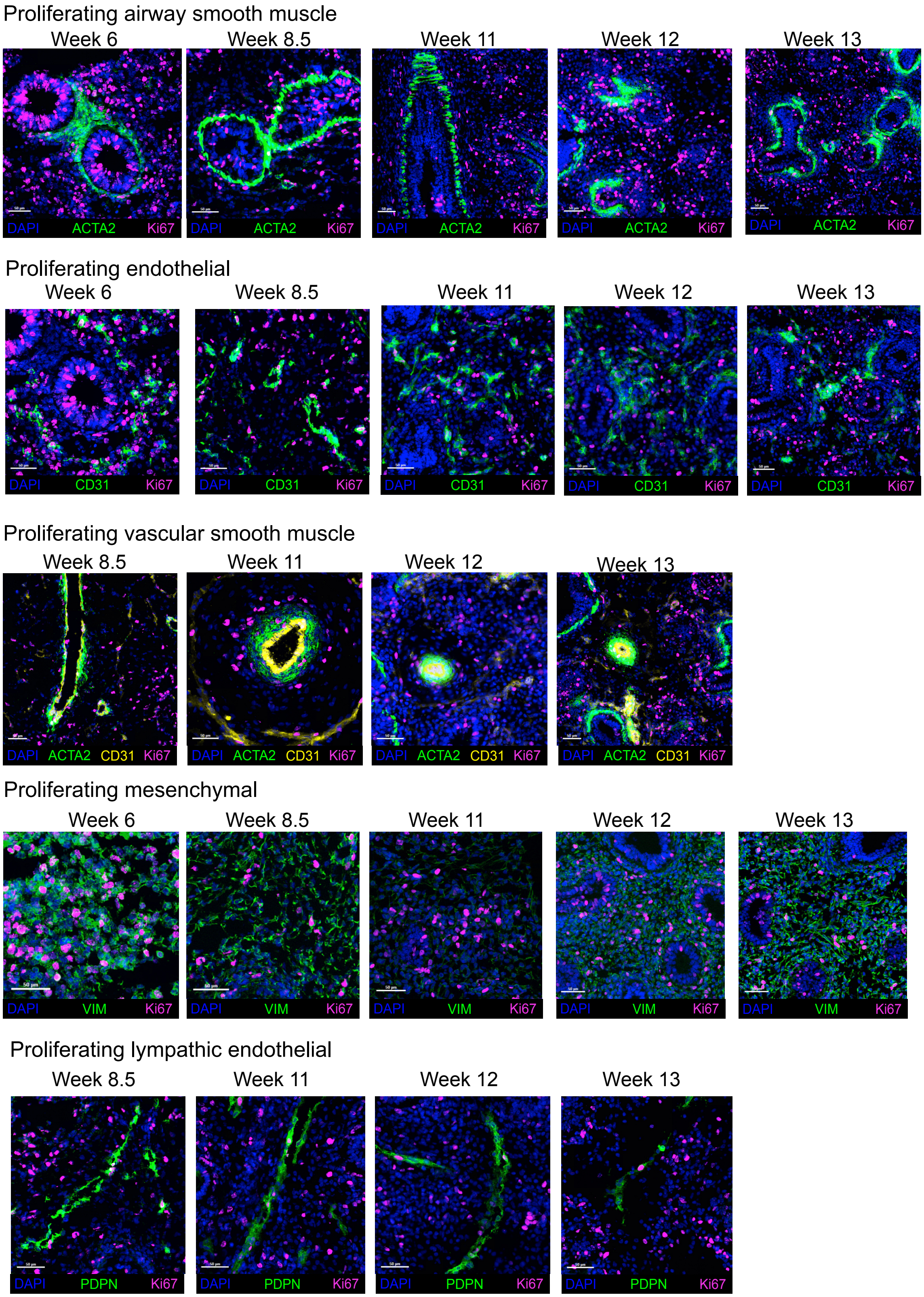
Proliferating airway smooth muscle
Week 6
Week 8.5
Week 11
Week 12
Week 13
DAPI ACTA2 Ki67
Proliferating endothelial
Week 6
Week 8.5
Week 11
Week 12
Week 13
DAPI CD31 Ki67
Proliferating vascular smooth muscle
Week 8.5
Week 11
Week 12
Week 13
DAPI ACTA2 CD31 Ki67
Proliferating mesenchymal
Week 6
Week 8.5
Week 11
Week 12
Week 13
DAPI VIM Ki67
Proliferating lymphatic endothelial
Week 8.5
Week 11
Week 12
Week 13
DAPI PDPN Ki67

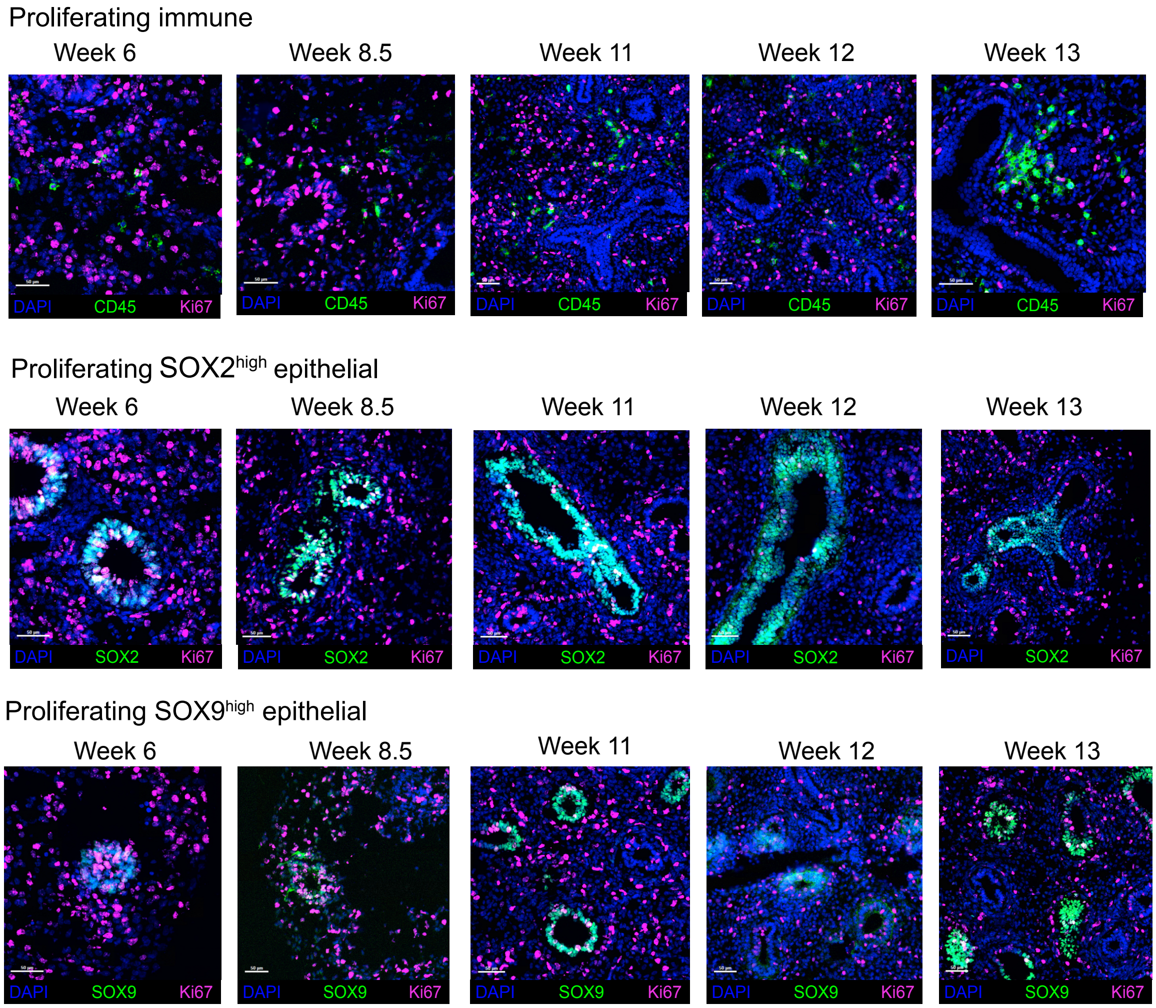


**Supplementary Figure 17. Expression patterns of Ki67 across the main cell types of the human developing lung.** Expression of Ki67 (magenta) was overlaid with expression of ACTA2 (green) for airway smooth muscle cells, CD31 (green) for endothelial cells, ACTA2 (green) and CD31 (yellow) for vascular smooth muscle cells, VIM (green) for mesenchymal cells, PDPN (green) for lymphatic endothelial cells, CD45 (green) for immune cells, SOX2 (green) for $SOX2^{high}$ epithelial cells and SOX9 (green) for $SOX9^{high}$ epithelial cells.

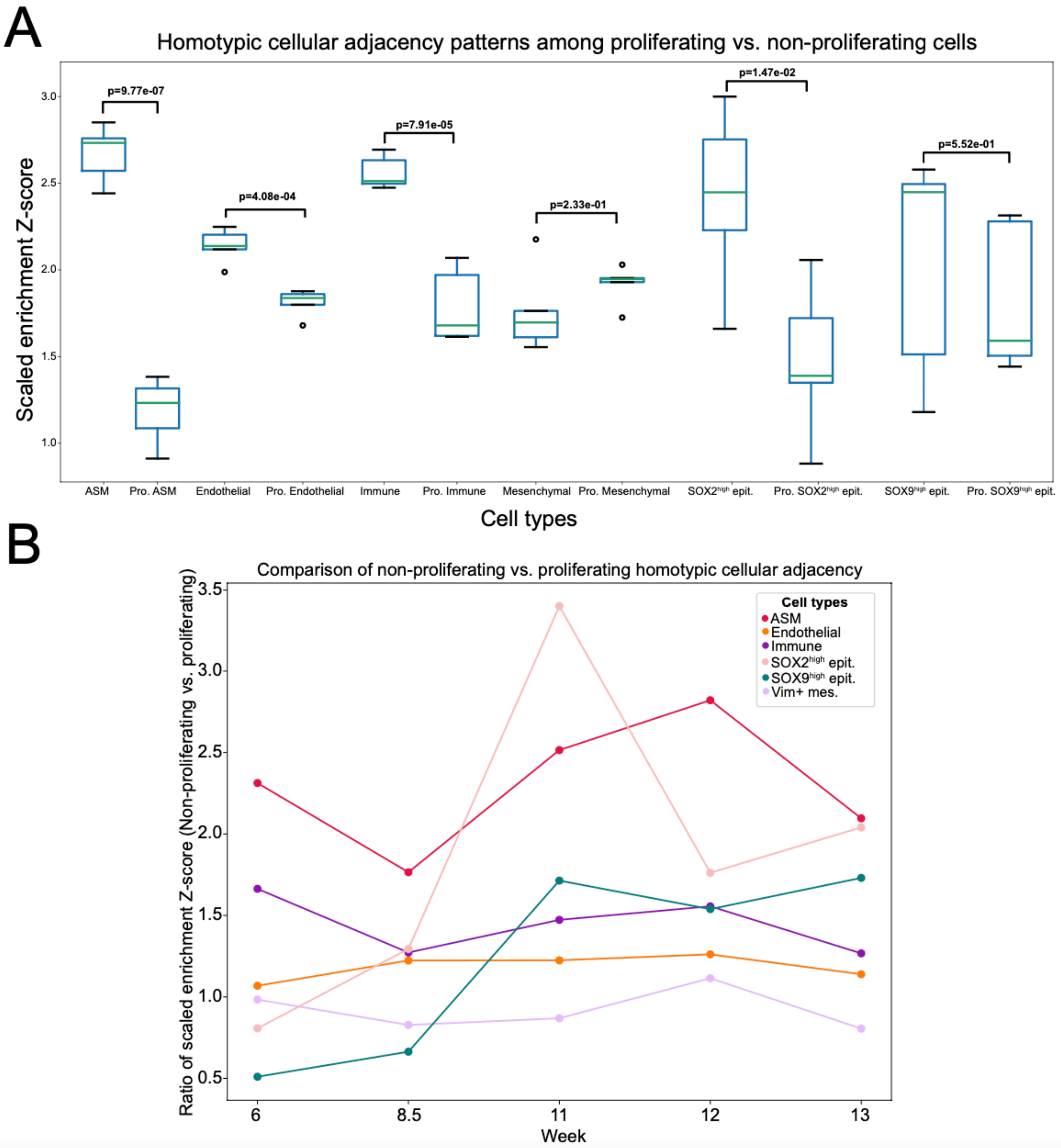


**Supplementary Figure 18. Comparison of homotypic cellular adjacency patterns among the non-proliferating cell types and their proliferating counterparts in the developing human lung. A)** Boxplots summarizing the scaled neighborhood enrichment Z-scores of six different cell types for its own cell type, where the neighborhood enrichment score for each non-proliferating and proliferating cell type across all the five developmental time points was compared in a two-sided Student's t-test. Source data for this figure is provided within the Source Data file. **B)** Lineplots summarizing the ratio of scaled enrichment Z-scores in non-proliferating versus proliferating counterparts of each cell type across the five developmental timepoints, where each line represents a different cell type. Source data for this figure is provided within the Source Data file.

**A**

Week 6

| Original Ann. | Simplified Ann. |
|---|---|
| Airway smooth muscle | Airway smooth muscle |
| Endothelial | Endothelial |
| Ki67+ mesenchymal | Mesenchymal |
| Mesothelium | Mesenchymal |
| Immune | Immune |
| $SOX2^{high}$ epithelial | $SOX2^{high}$ epithelial |
| $SOX9^{high}$ epithelial | $SOX9^{high}$ epithelial |
| Vim+ mesenchymal | Mesenchymal |

Week 8.5

| Original Ann. | Simplified Ann. |
|---|---|
| Airway smooth muscle | Airway smooth muscle |
| Endothelial | Endothelial |
| KI67+ mesenchymal | Mesenchymal |
| Lymphatic endothelial | Lymphatic endothelial |
| Immune | Immune |
| $SOX2^{high}$ epithelial | $SOX2^{high}$ epithelial |
| $SOX9^{high}$ epithelial | $SOX9^{high}$ epithelial |
| Vascular smooth muscle | Vascular smooth muscle |
| Vim+ mesenchymal | Mesenchymal |
| Neuronal | Mesenchymal |
| Chondroblasts | Mesenchymal |

Week 11

| Original Ann. | Simplified Ann. |
|---|---|
| Adventitial fibroblast | Mesenchymal |
| Airway fibroblast | Mesenchymal |
| Airway smooth muscle | Airway smooth muscle |
| Chondroblast | Mesenchymal |
| Endothelial | Endothelial |
| Ki67+ mesenchymal | Mesenchymal |
| Lymphatic endothelial | Lymphatic endothelial |
| Immune | Immune |
| Neuronal | Mesenchymal |
| $SOX2^{high}$ epithelial | $SOX2^{high}$ epithelial |
| $SOX9^{high}$ epithelial | $SOX9^{high}$ epithelial |
| Vascular smooth muscle | Vascular smooth muscle |
| Vim+ mesenchymal | Mesenchymal |

Week 12

| Original Ann. | Simplified Ann. |
|---|---|
| Airway smooth muscle | Airway smooth muscle |
| Endothelial | Endothelial |
| Ki67+ mesenchymal | Mesenchymal |
| Lymphatic endothelial | Lymphatic endothelial |
| Immune | Immune |
| $SOX2^{high}$ epithelial | $SOX2^{high}$ epithelial |
| $SOX9^{high}$ epithelial | $SOX9^{high}$ epithelial |
| Vascular smooth muscle | Vascular smooth muscle |
| Vim+ mesenchymal | Mesenchymal |
| Neuronal | Mesenchymal |

Week 13

| Original Ann. | Simplified Ann. |
|---|---|
| Airway smooth muscle | Airway smooth muscle |
| Chondroblast | Mesenchymal |
| Endothelial | Endothelial |
| Ki67+ mesenchymal | Mesenchymal |
| Lymphatic endothelial | Lymphatic endothelial |
| Neuronal | Mesenchymal |
| Immune | Immune |
| Pericytes | Mesenchymal |
| $SOX2^{high}$ epithelial | $SOX2^{high}$ epithelial |
| $SOX9^{high}$ epithelial | $SOX9^{high}$ epithelial |
| Vascular smooth muscle | Vascular smooth muscle |
| Vim+ mesenchymal | Mesenchymal |

**B**

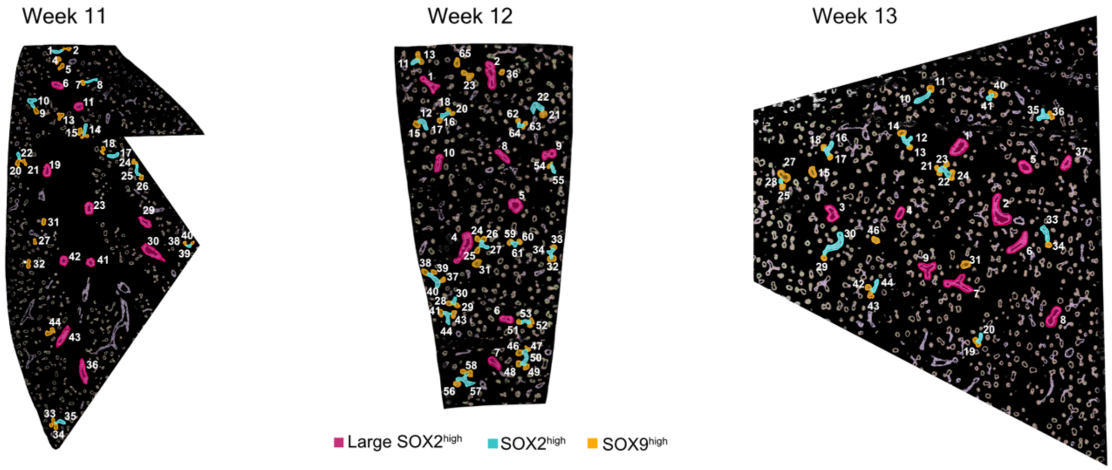


**C**

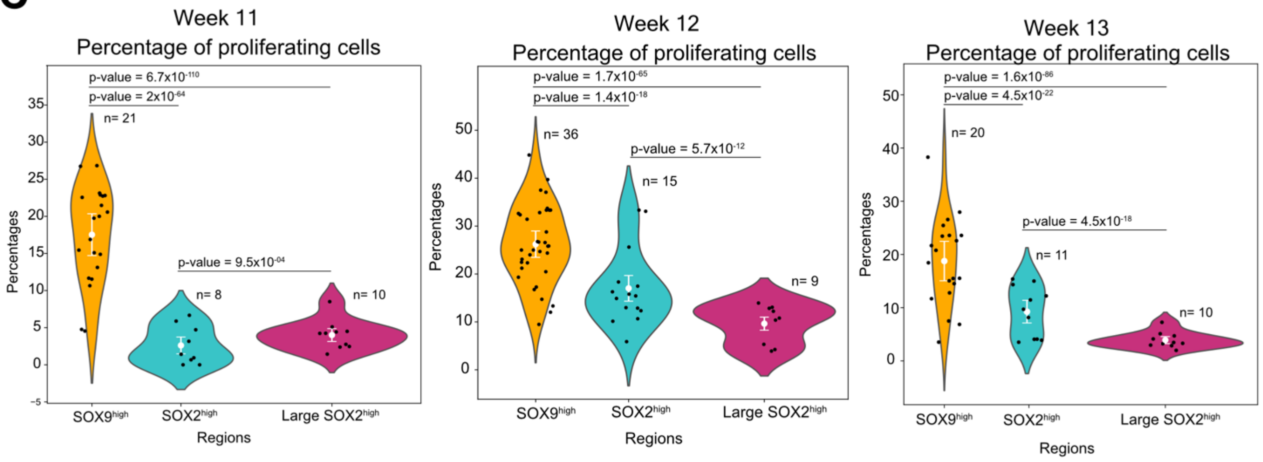

**Supplementary Figure 19. Cell-type specific proliferation patterns in the developing human lung. A)** Simplified annotations used for the proliferation analysis. The first column shows the original cell type annotation and the second column shows the simplified annotations for the proliferation analysis. **B)** The selected regions for analysis and comparison of proliferation patterns in relatively smaller $SOX2^{high}$ (cyan), relatively larger $SOX2^{high}$ (magenta) and $SOX9^{high}$ (orange) epithelial regions. ID of each identified and used region for analysis is stated on the tissue images. **C)** Violin plots displaying the distribution of the fraction of proliferating cells in relatively smaller $SOX2^{high}$ (cyan), relatively larger $SOX2^{high}$ (magenta) and $SOX9^{high}$ (orange) epithelial regions at each week. The x-axis represents the three different airway structures, and the y-axis indicates the percentage of proliferating cells in each of the (n) number of manually selected regions represented as a dot, and the error bar represents mean±2×SEM (standard error of the mean). The statistical significance of the differences in proliferation degree are summarized with a two-sided Fisher's exact test p-values. Source data for this figure is provided within the Source Data file.

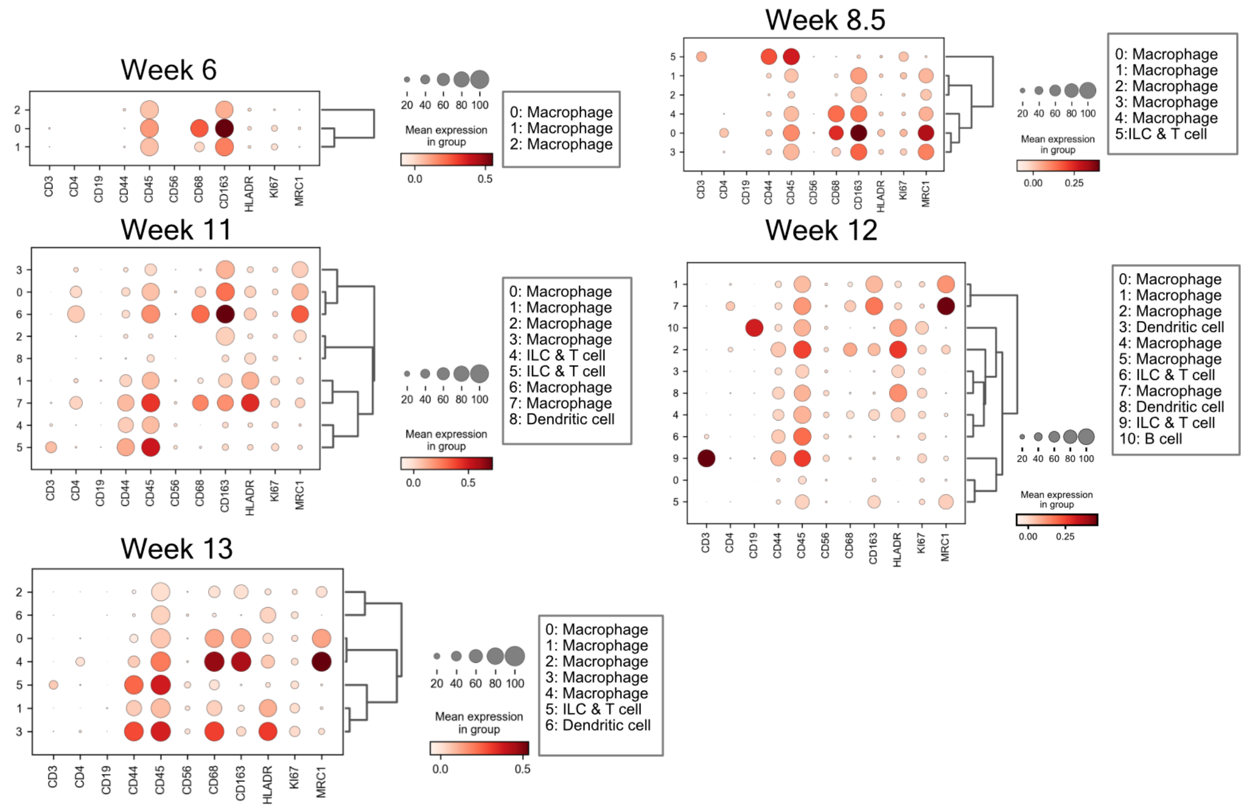


**Supplementary Figure 20. Subclustering of the major immune cell clusters identified in developing human lung.** The major immune cluster identified at each week was re-clustered and annotated based on the marker expressions in the dot plots. The size of the circles indicates the percentage of cells expressing the marker in each cluster, where dark red color indicates high and white color indicates low expression.

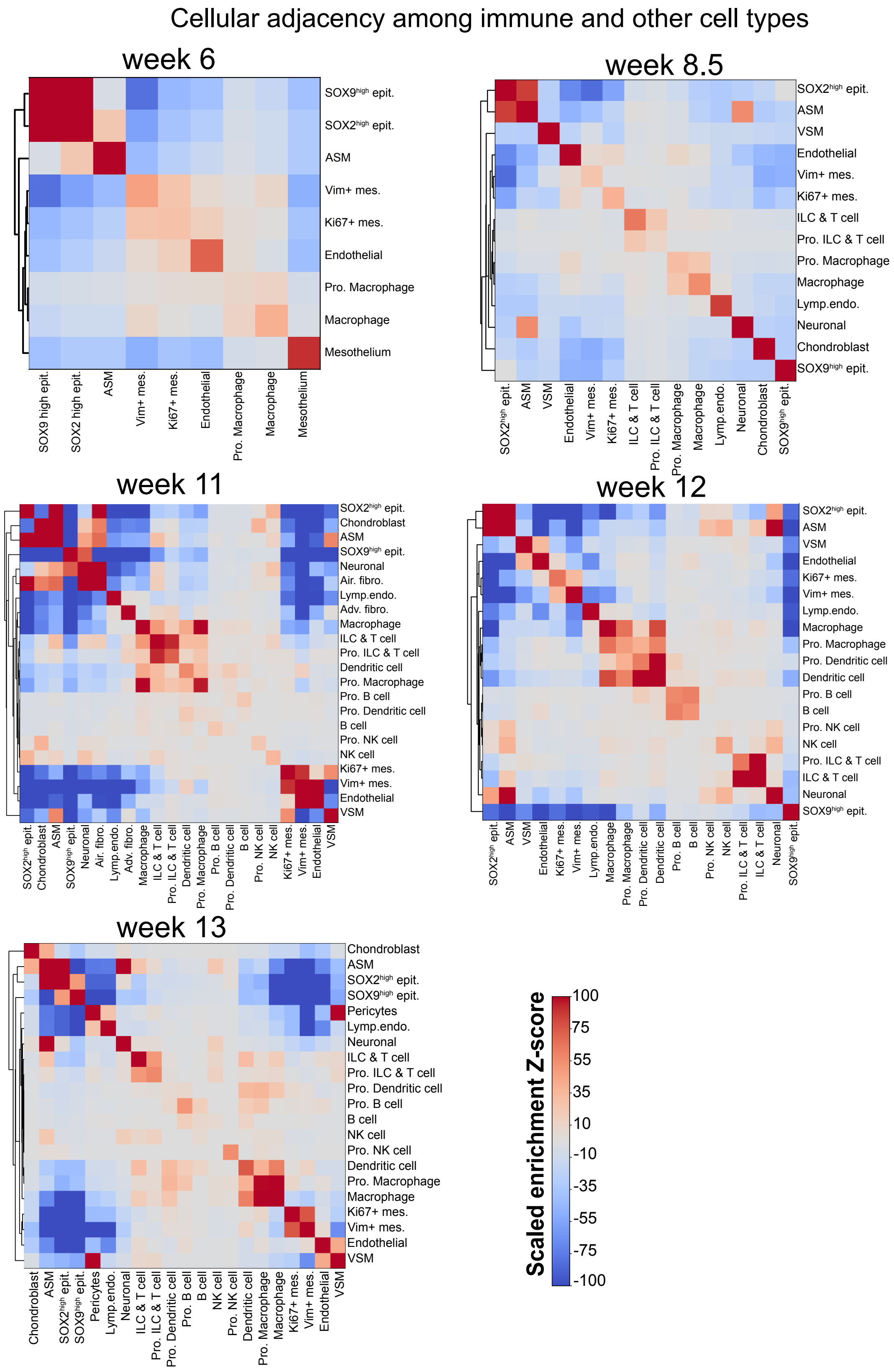


**Supplementary Figure 21. Heatmaps with dendrograms summarizing enrichment of cellular adjacency patterns among immune and other cell types based on the scaled Z-scores.** Dark red color indicates cellular adjacency patterns with high enrichment Z-score and dark blue color indicates cellular adjacency patterns with low enrichment Z-score.

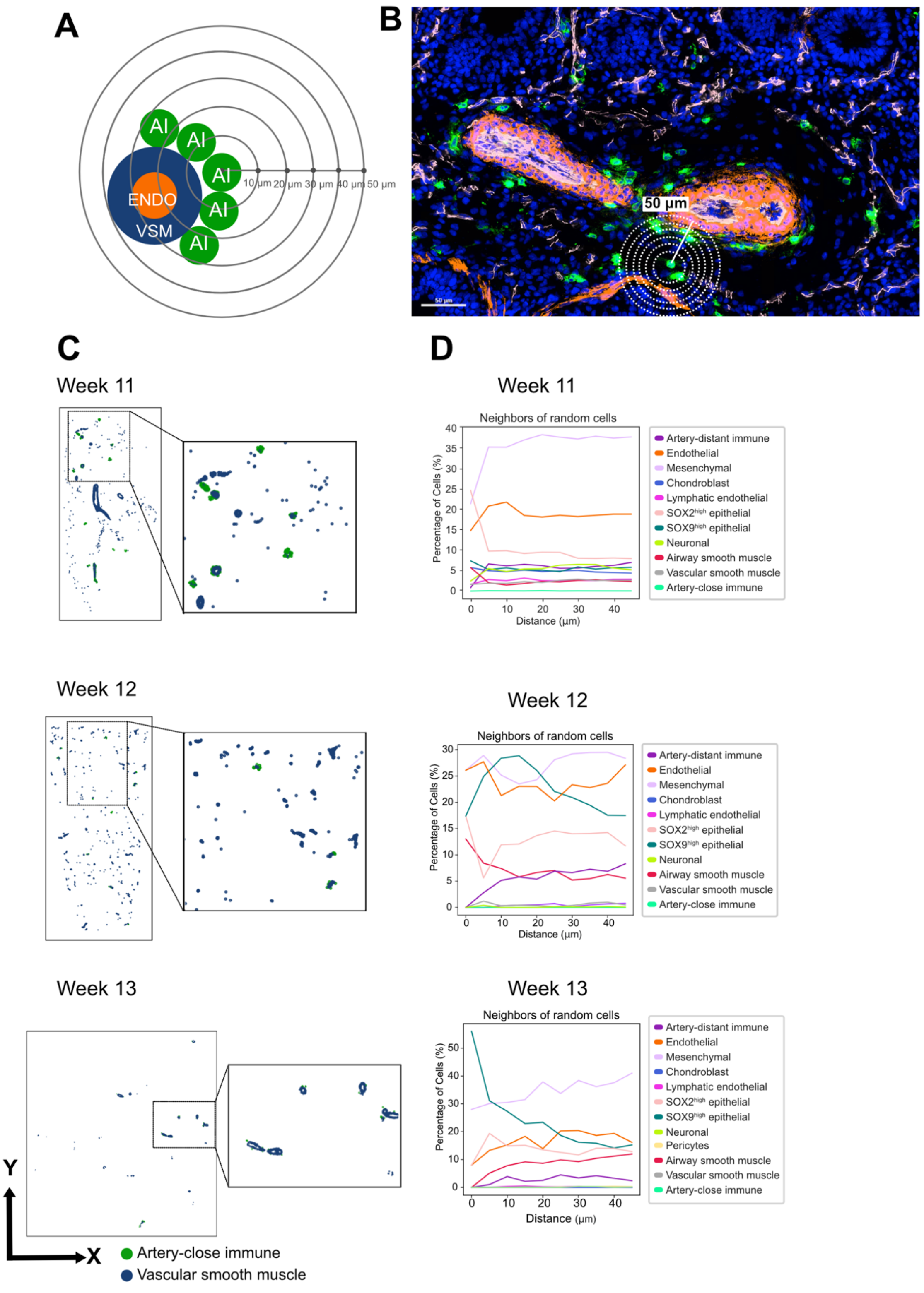
A
AI
ENDO
VSM
10 μm
20 μm
30 μm
40 μm
50 μm
B
50 μm
C
Week 11
Week 12
Week 13
Y
X
Artery-close immune
Vascular smooth muscle
D
Week 11
Neighbors of random cells
Percentage of Cells (%)
Distance (μm)
Artery-distant immune
Endothelial
Mesenchymal
Chondroblast
Lymphatic endothelial
SOX2high epithelial
SOX9high epithelial
Neuronal
Airway smooth muscle
Vascular smooth muscle
Artery-close immune
Week 12
Neighbors of random cells
Percentage of Cells (%)
Distance (μm)
Artery-distant immune
Endothelial
Mesenchymal
Chondroblast
Lymphatic endothelial
SOX2high epithelial
SOX9high epithelial
Neuronal
Airway smooth muscle
Vascular smooth muscle
Artery-close immune
Week 13
Neighbors of random cells
Percentage of Cells (%)
Distance (μm)
Artery-distant immune
Endothelial
Mesenchymal
Chondroblast
Lymphatic endothelial
SOX2high epithelial
SOX9high epithelial
Neuronal
Pericytes
Airway smooth muscle
Vascular smooth muscle
Artery-close immune

**Supplementary Figure 22. Neighborhood cellular composition analysis of immune cells encircling the arteries. A)** Scheme describing the neighborhood cellular composition analysis of the artery-close immune cells (AI) encircling vascular smooth muscle cells (VSM) and endothelial cells (ENDO). **B)** Snapshots of immune cells encircling arteries and a representation of the analyzed cellular composition for a radius of 50 µm around each detected artery-close immune cell. DAPI in blue, CD45 (immune marker) in green, ACTA2 (vascular smooth muscle marker) in orange and CD144 (endothelial marker) in pink. **C)** The xy-scatter plots show the location of immune cells detected around arteries and vascular smooth muscle cells. **D)** Line plots summarizing the cell type and count of neighbors of randomly selected cells within a radius of 50 µm for weeks 11,12 and 13. The x-axis represents the distance in µm and the y-axis represents the fraction of cells. Source data for this figure is provided within the Source Data file.

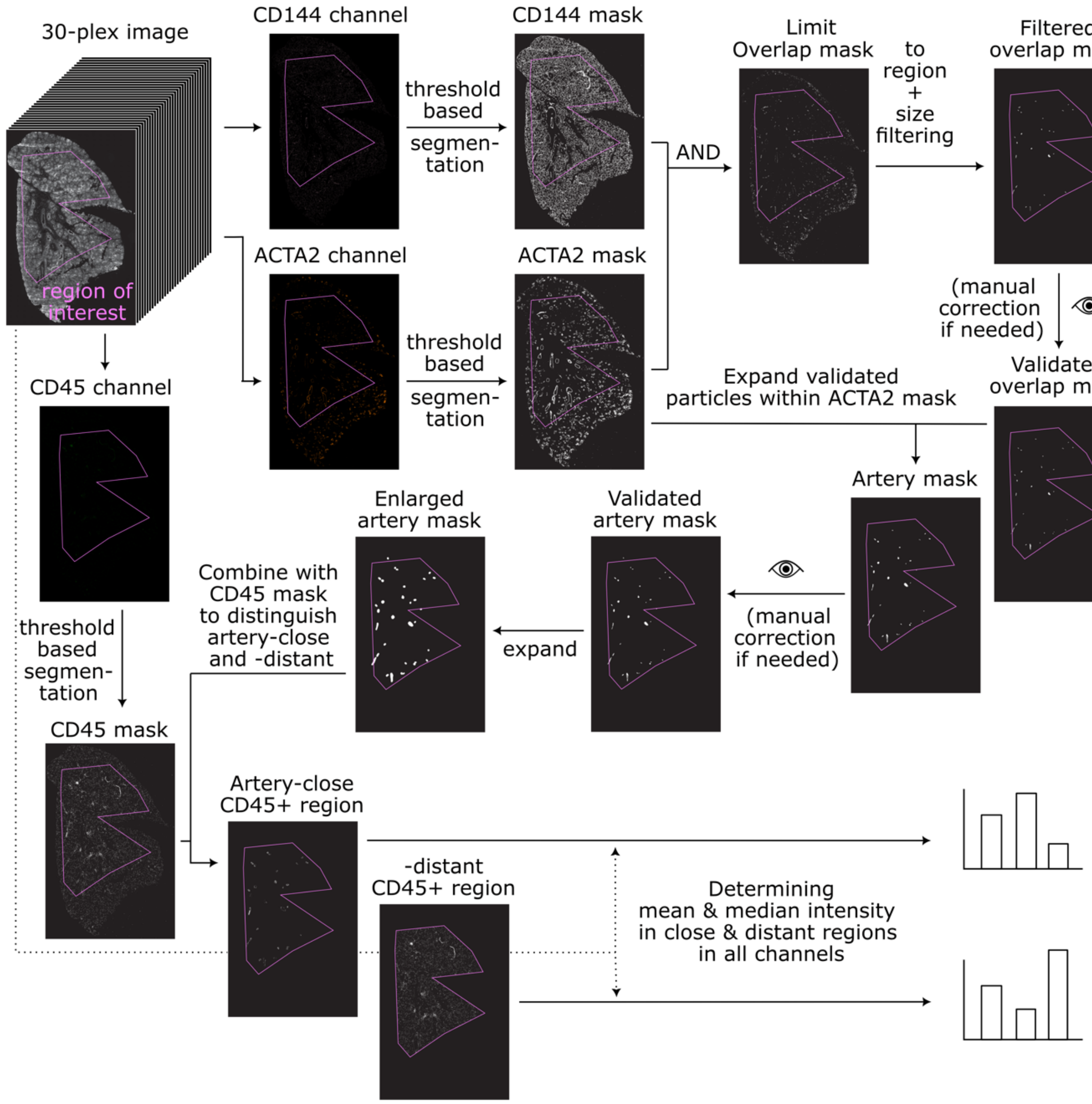


**Supplementary Figure 23. Image analysis workflow to validate marker expression patterns of artery-close and -distant immune cells without instance segmentation.** The workflow was developed as an ImageJ Macro available at https://github.com/CellProfiling/HDCA-FetalLung-SpatialProteomics/ (10.5281/zenodo.11650173). Briefly, the raw image stack was loaded and the user was prompted to select a region of interest, to which all analysis including threshold determination would be restricted. Next, the channel images for ACTA2, CD144, and CD45 were extracted, pre-processed, and subjected to semantic segmentation using ImageJ's Triangle threshold algorithm (for more details see Methods). Masks derived from ACTA2 and CD144 channel images were combined using the logical AND operation, and the resulting mask was filtered with a size filter: all objects smaller than 2000 px were removed. After this step, the mask was displayed to the user asking to validate it and to eventually add any missed-out artery regions or to remove artifacts. The objects in the validated mask were expanded within the ACTA2 mask to extend to the whole artery region. The user was asked again for validation. Then, the expanded validated mask was expanded again using a maximum filter (radius of 80 px = 40 µm) to cover the peri-arterial regions. This mask was then used to dissect the immune cell mask coming from the CD45 channel into artery-close and artery-distant regions. The artery-close and the -distant mask were used to determine the mean and median intensity within -close and -distant immune cell regions, in all image channels of the raw 30-plex image. The example lung tissue image shown in the workflow is from the 11-pcw-old sample.

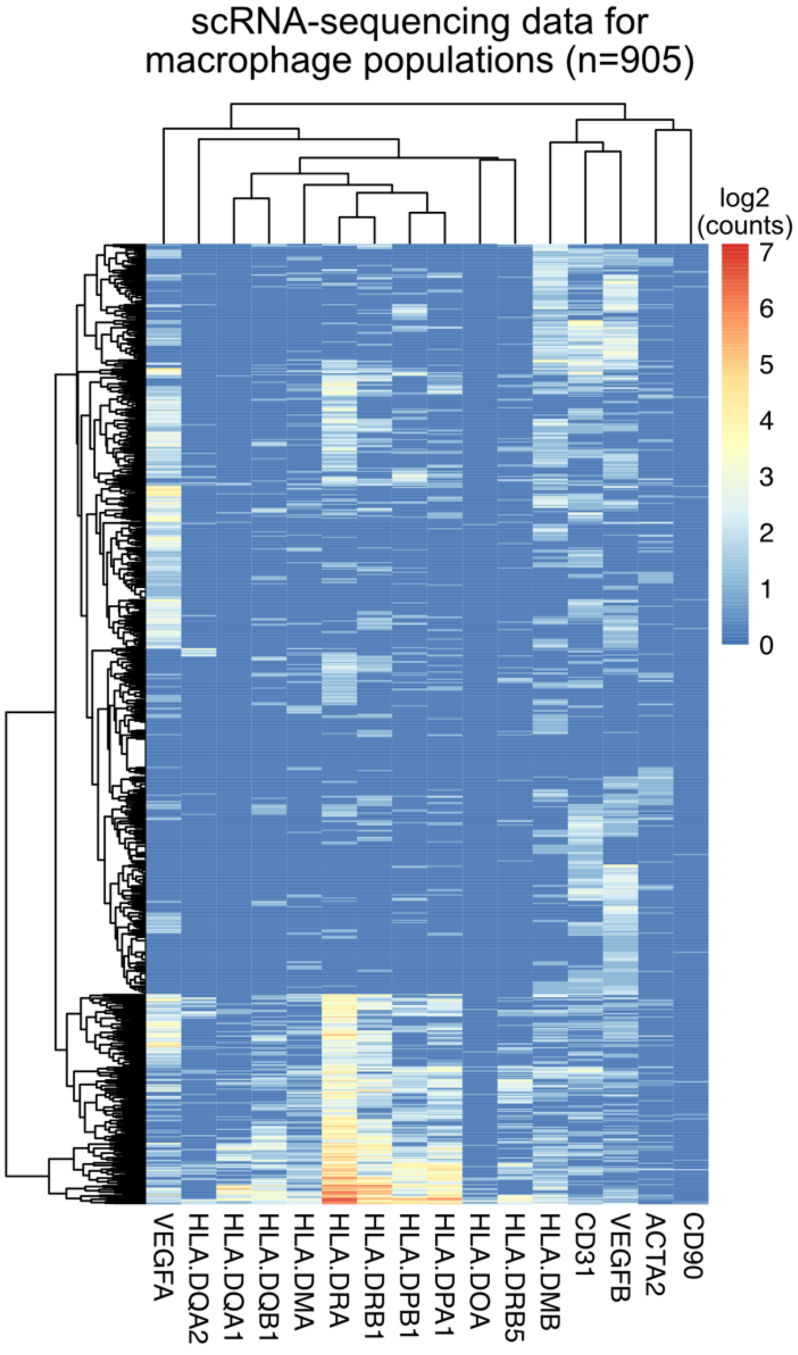


**Supplementary Figure 24. Heatmap summarizing the expression patterns of selected genes in macrophage populations which were identified in single-cell transcriptomics human developing lung dataset by Sountoulidis *et al*[1].** A total of 905 cells (on the y-axis) were annotated as macrophages across the developmental weeks from 5 to 14. The heatmap summarizes the log2(normalized UMI counts+1) for 16 selected genes.
