## Supplementary Table 1 for "High-parametric protein maps reveal the spatial organization in early-developing human lung"

### LIST OF ALL TESTED ANTIBODIES

| Antibodies | Catalog number | Clone | RRID | Target | Selection Reason | Barcodes of Akoya Immune Panel Antibodies | Regular IF Dilution | Regular IF Tested Tissue | In-House Conjugated/Barcode | Failed Step | Phenocycler Dilutions | Worked in the run |
|---|---|---|---|---|---|---|---|---|---|---|---|---|
| ACTA2 | ab119952 | 4A4 | AB_10901360 | Smooth muscle actin | sc-RNAseq | N/A | 1/200 | Fetal lung 13 pcw | BX028 | N/A | 1/70 | Yes |
| ASCL1 | ab240385 | EPR19840 | | Neuroendocrine cells | sc-RNAseq | N/A | 1/50- 1/100 | Fetal Lung 8.5 pcw | N/A | Failed in Regular IF | N/A | N/A |
| CD104 | 4250008 | 58XB4 | | Epithelial cells | Akoya Immune Panel | BX005 | N/A | N/A | N/A | Failed in the run | N/A | N/A |
| CD11c | 4350012 | S-HCL-3 | | Dendritic cells/ macrophages | Akoya Immune Panel | BX027 | N/A | N/A | N/A | Failed in the run | N/A | N/A |
| CD123 | # 14-1239-82 | 6H6 | AB_467453 | Endothelial cells&Hematopoetic stem cells | Literature/sc-RNAseq | N/A | 1/100 | Tonsil | BX54 | N/A | 1/100 | Yes |
| CD14 | # 14-0149-82 | 61D3 | | Monocyte | Literature/sc-RNAseq | N/A | 1/50 | Tonsil/Spleen | BX006 | Failed in the run | N/A | N/A |
| CD144 | # 14-1449-82 | 16B1 | AB_467495 | Endothelial cells&Hematopoetic stem cells | Literature/sc-RNAseq | N/A | 1/100 | Pancreas | BX016 | N/A | 1/100 | Yes |
| CD163 | HPA046404 | | | Macrophages | Literature/sc-RNAseq | N/A | 1/130 | Fetal lung 13 pcw | BX025 | Failed in Conjugation | N/A | N/A |
| CD163 | # MA5-17716 | GHI/61 | AB_2539106 | Macrophages | Literature/sc-RNAseq | N/A | 1/50 | Fetal lung 13 pcw | BX049/ BX005 | N/A | 1/50 | Yes |
| CD19 | 4350003 | HIB19 | | B cells | Akoya Immune Panel | BX003 | N/A | N/A | N/A | N/A | 1/100 | Yes |
| CD2 | 4250005 | RPA-2.10 | | T cells | Akoya Immune Panel | BX002 | N/A | N/A | N/A | Failed in the run | N/A | N/A |
| CD20 | # 14-0202-82 | L26 | | B cells | Literature/sc-RNAseq | N/A | 1/100- 1/1000 | Spleen | N/A | Failed in Regular IF | N/A | N/A |
| CD21 | 4150009 | Bu32 | | B cells | Akoya Immune Panel | BX013 | N/A | N/A | N/A | Failed in the run | N/A | N/A |
| CD278 | 4250013 | C398.4A | | Activated helper and cytotoxic cells | Akoya Immune Panel | BX017 | N/A | N/A | N/A | Failed in the run | N/A | N/A |
| CD279 | 4250010 | EH12.2H7 | | T cells | Akoya Immune Panel | BX014 | N/A | N/A | N/A | Failed in the run | N/A | N/A |
| CD3 | 4350008 | UCHT1 | | T cells | Akoya Immune Panel | BX015 | N/A | N/A | N/A | N/A | 1/200 | Yes |
| CD31 | 4250009 | WM59 | | Endothelial cells | Akoya Immune Panel | BX032 | N/A | N/A | N/A | N/A | 1/200 | Yes |
| CD34 | 4250020 | 561 | | Endothelial cells | Akoya Immune Panel | BX035 | N/A | N/A | N/A | N/A | 1/200 | Yes |
| CD38 | 4150007 | FF | | General immune cells marker | Akoya Immune Panel | BX007 | N/A | N/A | N/A | Failed in the run | N/A | N/A |
| CD4 | 4350010 | SK3 | | Helper T cells | Akoya Immune Panel | BX021 | N/A | N/A | N/A | N/A | 1/200 | Yes |
| CD41 | HPA031169 | | | Megakaryocytes | Literature/sc-RNAseq | N/A | 1/250 | Fetal lung 13 pcw | BX041 | Failed in Conjugation | N/A | N/A |
| CD44 | # MA4400 | Hermes-1 | AB_223517 | Hematopoetic stem cells | Literature/sc-RNAseq | N/A | 1/50 | Fetal Lung 12 pcw | BX020 | N/A | 1/100 | Yes |
| CD45 | 4150003 | HI30 | | General immune cells marker | Akoya Immune Panel | BX001 | N/A | N/A | N/A | N/A | 1/100 | Yes |
| CD56 | # MA1-06801 | 123C3 | | NK cells | Literature/sc-RNAseq | N/A | 1/100 | Pancreas | BX029 | Regular IF | N/A | N/A |
| CD56 | ab251595 | CAL53 | | NK cells | Literature/sc-RNAseq | N/A | 1/50 | Fetal lung 13 pcw | BX029 | N/A | 1/50 | Yes |
| CD59 | # MA5-17046 | 8D2B8 | | Lineage marker | Literature | N/A | 1/200 | Pancreas | BX033 | Failed in the run | N/A | N/A |
| CD68 | # 14-0688-82 | KP1 | AB_11151139 | Macrophage | Literature/sc-RNAseq | N/A | 1/100 | Tonsil | BX10 | N/A | 1/100 | Yes |
| CD69 | 4250022 | FN50 | | Lymphocyte activation marker | Akoya Immune Panel | BX041 | N/A | N/A | N/A | Failed in the run | N/A | N/A |
| CD7 | 4150022 | CD7-6B7 | | Differentiation marker for T cells | Akoya Immune Panel | BX025 | N/A | N/A | N/A | Failed in the run | N/A | N/A |
| CD8 | 4150004 | SK1 | | Cytotoxic T cells | Akoya Immune Panel | BX004 | N/A | N/A | N/A | Failed in the run | N/A | N/A |
| CD9 | 4150016 | HI9a | | Myeloid cell lineage cells | Akoya Immune Panel | BX028 | N/A | N/A | N/A | Failed in the run | N/A | N/A |
| CD90 | 4150021 | 5E10 | | Stem cells | Akoya Immune Panel | BX022 | N/A | N/A | N/A | N/A | 1/200 | Yes |
| CKIT | #348800 | 2B8 | | Hematopoetic stem cells | Literature/sc-RNAseq | N/A | 1/125 | Mouse cerebellum | BX45 | Failed in the run | N/A | N/A |
| CKIT | HPA073252 | | | Hematopoetic stem cells | Literature/sc-RNAseq | N/A | 1/30 | Fetal lung 13 pcw | BX050 | Failed in Conjugation | N/A | N/A |
| CLDN5 | # 35-2500 | 4C3C2 | AB_2533200 | Endothelial cells | sc-RNAseq | N/A | 1/100 | Fetal lung 7 w | BX041 | N/A | 1/100 | Yes |
| COL1A1 | MAB6220-100 | # 816161 | | Mesenchymal | sc-RNAseq | N/A | 1/100 | Fetal lung 13 pcw | BX002 | N/A | 1/300 | Yes |
| CSF1R | ab240265 | SP211 | | Macrophages | sc-RNAseq | N/A | 1/100 | Fetal lung 13 pcw | BX017/BX036 | Failed in the run | N/A | N/A |
| CXCR4 | # 35-8800 | 12G5 | | Hematopoetic stem cells | Literature/sc-RNAseq | N/A | 1/50-1/500 | Spleen | N/A | Failed in Regular IF | N/A | N/A |
| DCN | HPA003315 | | | Mesenchymal | sc-RNAseq | N/A | 1/200 | Fetal lung 13 pcw | BX013 | N/A | 1/50 | Yes |
| Ecadherin | 4250021 | 4A2C7 | | Epithelial Cells | Akoya FFPE Panel | BX014 | N/A | N/A | N/A | N/A | 1/200 | Yes |
| EPCAM | #14932682 | 1B7 | AB_795876 | Epithelial cells | sc-RNAseq | N/A | 1/100 | Fetal lung 13 pcw | BX042 | N/A | 1/400 | Yes |
| FLT3 | # PA5-34448 | | | Hematopoetic stem cells | Literature/sc-RNAseq | N/A | 1/50-1/500 | Spleen | N/A | Failed in Regular IF | N/A | N/A |
| GP1BA | HPA013316 | | | Megakaryocytes | Literature/sc-RNAseq | N/A | 1/110 | Fetal lung 13 pcw | BX045 | Failed in Conjugation | N/A | N/A |
| GPC3 | # MA5-17083 | 9C2 | | Alveolar fibroblast | sc-RNAseq | N/A | 1/100 | Fetal lung 7 w | N/A | Failed in Regular IF | N/A | N/A |
| GRHL2 | # MA5-31388 | CL3760 | | Mesenchyme | sc-RNAseq | N/A | 1/250 | Fetal lung 13 pcw | BX024 | Failed in Regular IF | N/A | N/A |
| HHIP | H00064399-M01 | 5D11 | | Lung airways | sc-RNAseq | N/A | 1/50 | Fetal lung 13 pcw | N/A | Failed in Regular IF | N/A | N/A |
| HLA-DR | 4250006 | L243 | | Antigen presenting cells | Akoya Immune Panel | BX026 | N/A | N/A | N/A | N/A | 1/200 | Yes |
| HPGD | NB200-179 | | | Aerocytes | sc-RNAseq | N/A | 1/100 | Fetal lung 7 w | N/A | Failed in Regular IF | N/A | N/A |
| IBA1 | ab221790 | EPR16589 | | Macrophages | sc-RNAseq | N/A | 1/50 | Fetal Lung 12 pcw | BX30, BX027 | Failed in Conjugation | N/A | N/A |
| IBA1 | ab220815 | EPR16588 | | Macrophages | sc-RNAseq | N/A | 1/100 | Fetal lung 7 w | BX036 | Failed in the run | N/A | N/A |
| Ki67 | 4250019 | B56 | | Proliferating cells | Akoya Immune Panel | BX047 | N/A | N/A | N/A | N/A | 1/200 | Yes |
| LYZ | ab185129 | EPR2994(2) | | Macrophages | sc-RNAseq | N/A | 1/1000 | Fetal lung 7 w | BX050 | Failed in Conjugation | N/A | N/A |
| MCM3 | SAB1404055 | 3E1 | | Proliferating cells | sc-RNAseq | N/A | 1/50 | Fetal lung 13 pcw | N/A | Failed in Regular IF | N/A | N/A |
| MRC1 | ab64693 | N/A | AB_1523910 | Macrophages | sc-RNAseq | N/A | 1/500 | Fetal lung 13 pcw | BX030 | N/A | 1/100 | Yes |
| NRXN1 | #703650 | 22H29L23 | | Cell adhesion | sc-RNAseq | N/A | 1/100 | Fetal lung 13 pcw | N/A | Failed in Regular IF | N/A | N/A |
| Pan-cytokeratin | 4150020 | AE-1/AE-3 | | Epithelial Cells | Akoya Immune Panel | BX019 | N/A | N/A | N/A | N/A | 1/300 | Yes |
| PDGFRB | MA535288 | ARC0009 | | Mesenchymal | sc-RNAseq | N/A | 1/100 | Fetal lung 13 pcw | N/A | Failed in Regular IF | N/A | N/A |
| PDGFRB | AF385 | | | Mesenchymal | sc-RNAseq | N/A | 1/100 | Fetal lung 13 pcw | N/A | Failed in Regular IF | N/A | N/A |
| Podoplanin | 4250004 | NC-08 | | Lymphatic endothelial cells | Akoya Immune Panel | BX023 | N/A | N/A | N/A | N/A | 1/200 | Yes |
| PRX | ab278083 | EPR24150-36 | | Capillaries | sc-RNAseq | N/A | 1/100 | Fetal lung 7 w | BX052 | N/A | 1/100 | Yes |
| PSAP | ab249391 | EPR10784(B) | | Macrophages | sc-RNAseq | N/A | 1/50 | Fetal lung 13 pcw | BX055 | Unsure Regular IF & Failed conjugation | N/A | N/A |
| RUNX1 | SAB1412471 | 3A1 | | Hematopoetic stem cells | Literature/sc-RNAseq | N/A | 1/50 | Fetal lung 13 pcw, Spleen | N/A | Failed in Regular IF | N/A | N/A |
| RUNX1 | H00000861-M06 | M06 | | Hematopoetic stem cells | Literature/sc-RNAseq | N/A | 1/300 | Fetal Lung 12 pcw | N/A | Failed in Regular IF | N/A | N/A |
| RUNX1 | HPA004176 | | | Hematopoetic stem cells | Literature/sc-RNAseq | N/A | 1/25 | Fetal lung 7 pcw | N/A | Failed in Regular IF | N/A | N/A |
| Serglycin | # PA5-50794 | | | Hematopoetic stem cells | Literature/sc-RNAseq | N/A | 1/100 | Fetal lung 13 pcw | BX025- BX030 | Failed in Conjugation | N/A | N/A |
| SFTPC | H00006440-M01 | 4A10 | | Type 2 alveolar cells | sc-RNAseq | N/A | 1/100 | Fetal lung 13 pcw | N/A | Failed in Regular IF | N/A | N/A |
| SOX2 | 14-9811-82 | Btjce | AB_11219471 | Proximal epithelial cells | sc-RNAseq | N/A | 1/250 | Fetal Lung 8.5 pcw | BX024 | N/A | 1/200 | Yes |
| SOX9 | MA5-17177 | 1B11 | | Distal epithelial cells | sc-RNAseq | N/A | 1/200 | Fetal Lung 8.5 pcw | N/A | Failed in Regular IF | N/A | N/A |
| SOX9 | AF3075 | | AB_2194160 | Distal epithelial cells | sc-RNAseq | N/A | 1/500 | Fetal lung 7 w | BX033 | N/A | 1/250 | Yes |
| TTF1 | ab216648 | EP1584Y | | Airway cells | sc-RNAseq | N/A | 1/250 | Whole embryo | BX007 | N/A | 1/100 | Yes |
| Vimentin | 677802 | O91D3 | AB_2565982 | Mesenchymal | sc-RNAseq | N/A | 1/50 | Fetal lung 13 pcw | BX045 | N/A | 1/300 | Yes |
| WT1 | # MA1-46028 | 6F-H2 | AB_962464 | Mesothelium | sc-RNAseq | N/A | 1/400 | Fetal lung 13 pcw | BX006 | N/A | 1/50 | Yes |

### LIST OF ANTIBODIES IN THE FINAL PANEL

| Antibodies | Catalog number | Clone | Target | Selection reason | Barcodes |
|---|---|---|---|---|---|
| ACTA2 | ab119952 | 4A4 | Smooth muscle actin | sc-RNAseq | BX028 |
| CD123 | # 14-1239-82 | 6H6 | Endothelial cells&Hematopoetic | Literature/sc-RNAseq | BX54 |
| CD144 | # 14-1449-82 | 16B1 | Endothelial cells&Hematopoetic | Literature/sc-RNAseq | BX016 |
| CD163 | # MA5-17716 | GHI/61 | Macrophages | Literature/sc-RNAseq | BX005 |
| CD19 | 4350003 | HIB19 | B cells | Akoya Immune Panel | BX003 |
| CD3 | 4350008 | UCHT1 | T cells | Akoya Immune Panel | BX015 |
| CD31 | 4250009 | WM59 | Endothelial cells | Akoya Immune Panel | BX032 |
| CD34 | 4250020 | 561 | Endothelial cells | Akoya Immune Panel | BX035 |
| CD4 | 4350010 | SK3 | Helper T cells | Akoya Immune Panel | BX021 |
| CD44 | # MA4400 | Hermes-1 | Hematopoetic stem cells | Literature/sc-RNAseq | BX020 |
| CD45 | 4150003 | HI30 | General immune cells marker | Akoya Immune Panel | BX001 |
| CD56 | ab251595 | CAL53 | NK cells | Literature/sc-RNAseq | BX029 |
| CD68 | # 14-0688-82 | KP1 | Macrophage | Literature/sc-RNAseq | BX10 |
| CD90 | 4150021 | 5E10 | Stem cells | Akoya Immune Panel | BX022 |
| CLDN5 | # 35-2500 | 4C3C2 | Endothelial cells | sc-RNAseq | BX041 |
| COL1A1 | MAB6220-100 | # 816161 | Mesenchymal | sc-RNAseq | BX002 |
| DCN | HPA003315 |  | Mesenchymal | sc-RNAseq | BX013 |
| Ecadherin | 4250021 | 4A2C7 | Epithelial Cells | Akoya FFPE Panel | BX014 |
| EPCAM | #14932682 | 1B7 | Epithelial cells | sc-RNAseq | BX042 |
| HLA-DR | 4250006 | L243 | Antigen presenting cells | Akoya Immune Panel | BX026 |
| Ki67 | 4250019 | B56 | Proliferating cells | Akoya Immune Panel | BX047 |
| MRC1 | ab64693 | N/A | Macrophages | sc-RNAseq | BX030 |
| Pan-cytokeratin | 4150020 | AE-1/AE-3 | Epithelial Cells | Akoya Immune Panel | BX019 |
| Podoplanin | 4250004 | NC-08 | Lymphatic endothelial cells | Akoya Immune Panel | BX023 |
| PRX | ab278083 | EPR24150-36 | Capillaries | sc-RNAseq | BX052 |
| SOX2 | 14-9811-82 | Btjce | Proximal epithelial cells | sc-RNAseq | BX024 |
| SOX9 | AF3075 |  | Distal epithelial cells | sc-RNAseq | BX033 |
| TTF1 | ab216648 | EP1584Y | Airway cells | sc-RNAseq | BX007 |
| Vimentin | 677802 | O91D3 | Mesenchymal | sc-RNAseq | BX045 |
| WT1 | # MA1-46028 | 6F-H2 | Mesothelium | sc-RNAseq | BX006 |
