## Supplementary Table 2 for "High-parametric protein maps reveal the spatial organization in early-developing human lung"

| | 6 pcw | 8.5 pcw | 11 pcw | 12 pcw | 13 pcw |
|---|---|---|---|---|---|
| **ACTA2** | 0.4576 | 13.494 | 0.8668 | 0.06 | 0.4301 |
| **CD3** | 3.3022 | 0.385 | 8.1543 | 3.2307 | 1.3552 |
| **CD4** | 23.0351 | 2.378 | 0.5302 | 6.6352 | 3.1262 |
| **CD19** | 1.9756 | 15.017 | 0.0528 | 1.8601 | 3.2164 |
| **CD31** | 0.1232 | 0.791 | 1.4586 | 5.507 | 8.179 |
| **CD34** | 0.0418 | 0.152 | 0.2816 | 0.461 | 3.198 |
| **CD44** | 3.3792 | 4.9 | 4.908 | 4.4517 | 24.0 |
| **CD45** | 1.5235 | 1.873 | 4.2471 | 3.4507 | 0.0737 |
| **CD56** | 0.1518 | 2.386 | 2.2077 | 0.0517 | 0.5819 |
| **CD68** | 20.372 | 14.277 | 20.3258 | 17.8244 | 9.886 |
| **CD90** | 10.0507 | 7.938 | 7.287 | 15.1866 | 7.0 |
| **CD123** | 0.8 | 0.230 | 2.3969 | 1.6126 | 0.1628 |
| **CD144** | 4.8 | 4.719 | 3.5992 | 6.6 | 1.782 |
| **CD163** | 2.1252 | 0.773 | 0.3883 | 5.6958 | 4.5078 |
| **CLDN5** | 6.3481 | 11.708 | 0.3179 | 0.2343 | 0.6094 |
| **COL1A1** | 3.105 | 3.083 | 0.8679 | 1.3849 | 1.0912 |
| **DCN** | 6.0 | 10.0 | 10.0 | 6.5 | 8.9 |
| **Ecadherin** | 0.3223 | 0.068 | 0.2211 | 0.9812 | 0.5115 |
| **EPCAM** | 1.9305 | 2.750 | 7.5438 | 6.864 | 10.9747 |
| **HLADR** | 0.3509 | 0.114 | 1.2001 | 5.124 | 1.3376 |
| **KI67** | 8.0 | 1.4 | 0.1298 | 0.0671 | 0.143 |
| **MRC1** | 4.8939 | 0.286 | 0.1221 | 15.609 | 2.2374 |
| **Pancytokeratin** | 0.7854 | 2.192 | 3.0316 | 0.2992 | 0.022 |
| **Podoplanin** | 0.8602 | 11.080 | 3.6212 | 1.5257 | 3.4892 |
| **PRX** | 0.055 | 0.025 | 0.143 | 0.0209 | 0.0209 |
| **SOX2** | 1.0604 | 0.232 | 0.4125 | 0.0363 | 3.0932 |
| **SOX9** | 7.9354 | 16.633 | 0.5709 | 4.0546 | 1.2672 |
| **TTF1** | 1.8656 | 5.008 | 6.1809 | 1.9426 | 0.198 |
| **Vimentin** | 8.9419 | 8.020 | 0.9834 | 4.4363 | 3.9633 |
| **WT1** | 13.0 | 9.0 | 3.5 | 8.437 | 0.4741 |
